# Bioassembly of region-specific fibrocartilage microtissues to engineer zonally defined meniscal grafts

**DOI:** 10.1101/2025.04.25.650568

**Authors:** Gabriela S. Kronemberger, Kaoutar Chattahy, Francesca D. Spagnuolo, Aliaa S. Karam, Daniel J. Kelly

## Abstract

Meniscal injuries are common orthopaedic problems which can impair knee function and lead to the development of osteoarthritis. While recent advances in tissue engineering have enabled the fabrication of meniscus-like grafts, these constructs do not fully replicate the zonal structure and composition of the native meniscus. In this study, we used fibrocartilage microtissues generated from meniscus progenitor cells (MPCs) as biological building blocks to biofabricate zonally defined meniscal grafts. MPCs isolated from the inner (iMPC) and outer (oMPC) regions of caprine menisci were used to engineer region-specific meniscal microtissues in a medium-high throughput manner. Both iMPC and oMPC derived microtissues were rich in glycosaminoglycans (GAGs) and collagen, with iMPC microtissues staining more intensely for type II collagen. These microtissues were assembled into two different physically confining moulds (cylindrical and ring-shaped), where they rapidly fused and generated a fibrocartilaginous graft over six weeks of culture. Both iMPC and oMPC assembled microtissues were rich in sGAG and type I collagen, however only the iMPC assembled microtissues stained strongly for type II collagen. We then explored the impact of the catabolic enzyme chondroitinase-ABC (cABC) on the composition and structural organization of the meniscal grafts. This temporal enzymatic treatment increased collagen fiber thickness without altering tissue phenotype. Finally, iMPC and oMPC microtissues were spatially assembled to biofabricate a scaled-up and zonally defined meniscal graft. This graft was phenotypically similar to the native meniscus, with all regions rich in collagen type I and an inner core rich in collagen type II and sGAG. These findings support the use of MPC derived microtissues as biological building blocks for the engineering of zonally defined meniscal grafts.

**Table of Content:**

- Zonally defined meniscus grafts were biofabricated using inner and outer meniscus progenitor cells microtissues.
- Region specific microtissues showed phenotypes similar to the native meniscus and successfully fused into cohesive constructs rich in sGAG and collagen.
- cABC treatment modulated collagen fiber formation and organization in assembled grafts.
- The assembled grafts maintained shape fidelity and may offer a promising strategy for meniscal repair.

Graphical Abstract:
Biofabrication of zonally defined meniscal grafts using meniscus progenitor cell (MPC)-derived microtissues. (A) Meniscus progenitor cells (MPCs) derived from the inner (iMPC), and outer (oMPC) regions of the menisci were aggregated into microtissues by cellular self-assembly process using a high-throughput agarose microwell system developed in-house. (B) The iMPC and oMPC microtissues were then separately fused in cylindrical and ring-shaped moulds to form region-specific assembled microtissues. (C) To enhance structural organization, chondroitinase ABC (cABC) enzymatic treatment was applied to the ring-shaped assembled microtissues. (D) Finally, the matured iMPC and oMPC assembled microtissues were assembled to create a scaled-up and zonally defined meniscal graft.

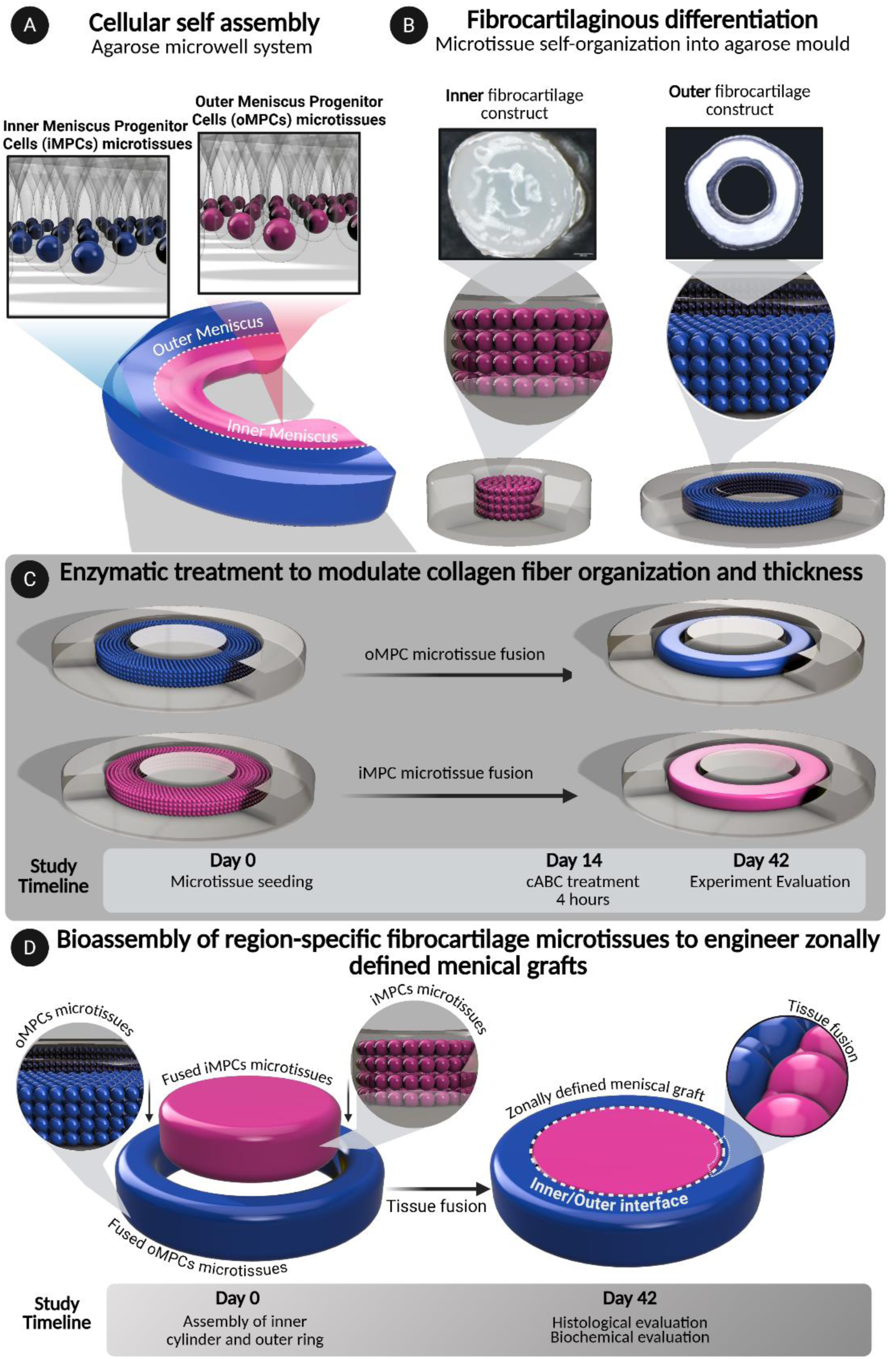

## 1. INTRODUCTION

The meniscus is a crescent-shaped fibrocartilage structure located on the medial and lateral aspects of the knee which plays a crucial role in load transmission and joint stability [1]. The meniscus is characterized by two anatomically distinct regions: the inner and outer zones. The extracellular matrix of the outer region is rich in type I collagen, while the avascular inner region contains higher levels of type II collagen and sulphated glycosaminoglycans (GAGs) [2,3]. Meniscal injuries are amongst the most frequent knee pathologies and can significantly increase the risk of developing osteoarthritis [4,5]. Current clinical treatments, such as meniscectomy, provide short term symptom relief but dramatically reduce meniscal functionality and accelerate the progression of osteoarthritis [6]. Alternative approaches, including meniscal allograft transplantation, synthetic meniscal substitutes such as Actifit®, and autologous chondrocyte implantation (ACI), have all been reported as treatment options for meniscal injuries, [6,7,8] however, long-term outcomes remain poor. These challenges have motivated increased interest in the field of meniscus tissue engineering [9]. While the use of emerging technologies such as controlled growth factor delivery, [10] enzymatic treatments, [11,12] novel biomaterials, [10] 3D bioprinting [13,14,15] and bioreactors for biophysical stimulation [16,17] have enabled the engineering of more complex meniscal constructs, these strategies typically fail to generate grafts that recapitulate the zonal structure and composition of the native tissue.

Bottom-up tissue engineering strategies aim to create complex constructs that mimic the structure and function of native tissues, often using multiple self-organised cellular units such as cellular aggregates, microtissues and organoids as biological building-blocks to generate scaled-up grafts [18,19]. Numerous studies have explored the potential of such microtissues to engineer musculoskeletal tissues such as bone [20,21,22,23] and cartilage [24,25,26,27,28]. Such approaches are particularly attractive for the engineering of spatially heterogeneous tissues like the meniscus, where phenotypically distinct microtissues can be engineered separately and then assembled to generate larger, spatially complex tissues or organs. Realising such an approach for meniscus tissue engineering requires the identification of a stem or progenitor cell populations capable of generating zone-specific meniscal microtissues. It has recently been demonstrated that the meniscus contains a population of progenitor cells with the capacity to generate region specific meniscal tissue [29,30,31]. Such meniscus progenitor cells (MPCs) represent an attractive potential cell source for the generation of region-specific meniscal microtissues.

Another key challenge in the field of cartilage and meniscus tissue engineering is the generation of grafts with a collagen content and organization similar to the native tissue. This can potentially be addressed using the temporal application of extracellular matrix (re)modelling enzymes such as chondroitinase ABC (cABC) and lysyl-oxidase (LOX) in culture, [11,32,33] which have been shown to modulate tissue composition and improve collagen organisation [34,35,36]. Furthermore, we have previously shown that cABC treatment enhanced the (re)modelling of hyaline cartilage grafts engineered via the fusion of articular cartilage microtissues, leading to the generation of more hierarchical organized [37]. Such enzymatic treatments could also potentially be used to improve the functionality of meniscal grafts engineered via the fusion of multiple fibrocartilage microtissues.

The goal of this study was to biofabricate zonally defined meniscus grafts using zone-specific fibrocartilage microtissues generated using MPCs. To this end MPCs from the inner (iMPC) and outer (oMPC) regions of the menisci were isolated to engineer region-specific meniscal microtissues in a medium-high throughput manner. iMPC and oMPC derived microtissues were then assembled in different spatial configurations to assess their capacity to fuse and develop region specific meniscal tissue. We then explored the impact of cABC treatment on the composition, structural organization, and overall collagen levels of the meniscal grafts. Finally, iMPC and oMPC microtissues were fused in a single construct aiming to biofabricate a cohesive, yet zonally defined, meniscus graft.

## 2. MATERIAL AND METHODS

### 2.1 Bone marrow mesenchymal stem/stromal cells (MSCs) isolation and expansion

MSCs were isolated from the sternum of skeletally mature, female, Saanen goats as described previously (18-19). Briefly, after the bone marrow pieces were harvested, they were gently cut into small pieces using a 10A scalpel. Next, the marrow pieces obtained were vortexed for 5 min in expansion medium (X-Pan) made from: Dulbecco’s Modified Eagle Medium (DMEM) + 100 U/mL penicillin + 100 μg/mL streptomycin (Gibco) + 10% (w/v) Fetal Bovine Serum (FBS) + Basic Fibroblastic Growth Factor 2 (FGF-2), to aid in liberating the cellular components. Then, the culture medium containing the cell suspension was aspirated and passed through a 40 μm cell strainer prior to counting and plating at a density of 5.7 × 10^4^ cells/cm^2^ and expanded under physioxic conditions (37° C in a humidified atmosphere with 5% CO_2_ and 5% O_2_) for improved chondrogenic differentiation. When the confluency in the flasks was at 80%, MSCs were trypsinized using 0.25% (w/v) trypsin Ethylenediaminetetraacetic acid (EDTA). For microtissues culture, MSCs were expanded from an initial density of 5000 cells/cm^2^ in X-Pan medium under physioxic conditions until passage 3.

### 2.2 Meniscus cell isolation and fibronectin selection

Medial and lateral menisci were obtained from skeletally mature, female, Saanen goats. Under sterile conditions, the menisci were rinsed twice with Dulbecco’s phosphate buffer (PBS) containing 100 U/ml penicillin and 100 µg/ml streptomycin (P/S) (Sigma-Aldrich, Dublin, Ireland). The tissue samples were then incubated in pronase (500 U/ml, 30 min at 37 °C) after which the menisci were dissected longitudinally into two parts: the inner zone and outer zones. Each zone was separately minced into 1-2 mm^3^ pieces and digested with collagenase type I solution (900 U/ml, 5 h at 37 °C) (Gibco). Following digestion, the solution was filtered through a 40 µm porous membrane, centrifuged, and resuspended in XPAN medium, consisting of DMEM GlutaMAX supplemented with 10% v/v FBS, 100 U/ml penicillin, and 100 µg/ml of streptomycin. Cells extracted from the inner and outer meniscal zones were cultured under two conditions: The first aimed to isolate a mixed population of inner (iFCs) and outer (oFCs) fibrochondrocytes, expanded at a high cell density (∼6000 cells/cm^2^). The second focused on isolating progenitor cells from the inner (iMPC) and outer (oMPC) meniscal region, expanded at a lower density (∼300 cells/cm^2^). These cells were subjected to a fibronectin selective adhesion, as previously described [14]. Briefly, petri dishes were coated with 10 μg/ml fibronectin in PBS containing 1 mM MgCl_2_, and 1 mM CaCl_2,_ and incubated overnight at 4°C. Cells from each region were seeded and incubated for 20 min at 37°C. Non-adherent cells were removed, and a fresh XPAN medium with 5 ng/ml FGF-2 (PeproTech, UK) was added. The adherent cells were then cultured at 37°C and 5% CO_2_.

### 2.3 Fibrochondrogenic differentiation

The fibrochondrogenic potential of progenitor groups was assessed at passages 2 and 4 using a 3D pellet culture protocol (Barcelo et al., 2023). Briefly, 2.5 × 10^5^ cells were centrifuged into pellets in chondrogenic media (CDM+), consisting of hgDMEM supplemented with (100 U/mL) penicillin, streptomycin (100 µg/mL)(both Gibco), sodium pyruvate (100 µg/mL), L-proline (40 µg/mL), L-ascorbic acid 2-phosphate (50 µg/mL), linoleic acid (4.7 µg/mL), bovine serum albumin (1.5 mg/mL), Insulin-Transferrin-Selenium, dexamethasone (100 nM), Amphotericin B (2.5 µg/mL) (all from Sigma), and human transforming growth factor-β3 (TGF-β) (10 ng/mL) (Peprotech). After 21 days in physioxic (37°C, 5% CO₂, 5% O₂), three pellets per cell region were used for biochemical quantification of DNA, sGAG, and collagen, while two were fixed in 4% paraformaldehyde (PFA) for histological analysis.

### 2.4 Osteogenic and adipogenic differentiation

Cells from the meniscus (iMPC, oMPC, iFC, and oFC) were seeded at ∼1×10^3^ cells/cm^2^ into 6-well plates and cultured in XPAN medium for 2-3 days. After reaching an 80% confluency, the basic medium was replaced with osteogenic medium [XPAN + 100 nM dexamethasone, 10 mM *β*-glycerophosphate and 0.05 mM ascorbic acid (all from Sigma)] and cultured for 21 days with media refreshed twice per week. For controls, cells were maintained in XPAN for the duration of culture. On day 21, cells were washed twice with PBS, fixed in iced ethanol at RT for 10 min, washed again with PBS and distilled water. Alizarin Red (1%, Sigma) was added for 1 min, followed by a final rinse with distilled water before phase contrast imaging (Zeiss) to investigate calcium deposition.

For adipogenic differentiation, cells were seeded at the same density as previously described for the osteogenic differentiation. The adipogenic medium was supplemented with 100 nM dexamethasone, 0.5 mM IBMX, and 50 μM indomethacin (all from Sigma). After 21 days in culture, cells were washed with PBS, incubated in 0.6% Oil Red O (ORO) solution at RT for 30 min, and washed with PBS until the background was clear. Positive adipogenic differentiation was confirmed by Oil red staining covering over 30% of the monolayer, evaluated by phase contrast microscopy (Zeiss).

### 2.5 Microtissue biofabrication and culture

iMPC, oMPC and MSC derived microtissues were fabricated by using an in-house high-throughput non-adherent agarose hydrogel microwell system as described previously (14-15). In this system, it is possible to fabricate up to 1,889 microtissues at once. For microtissue formation using a 400 microwells/mould, cells at passage 3 were seeded on top of each mould at final densities of 1 × 10³, 2 × 10³, or 4 × 10³ cells/microwell. For microtissue formation using an 1,889 microwells/mould, a density up to 3 x 10^3^ cells/microtissue was seeded in X-Pan media. After 24h, in order to allow cell aggregation, the microtissues were maintained in chondrogenic induced culture conditions consisting of hgDMEM GlutaMAX supplemented with 100 U/mL penicillin, 100 μg/mL streptomycin (both Gibco), 100 μg/mL sodium pyruvate, 40 μg/mL L-proline, 50 μg/mL L-ascorbic acid-2-phosphate, 4.7 μg/mL linoleic acid, 1.5 mg/mL bovine serum albumin, 1 × insulin–transferrin–selenium (ITS), 100 nM dexamethasone (all from Sigma), 2.5 μg/mL amphotericin B and 10 ng/mL of human transforming growth factor - beta 3 (TGF-β) (Peprotech, UK). The microtissues were cultured under physioxic conditions (37◦ C in a humidified atmosphere with 5% CO_2_ and 5% O_2_) for up to 3 weeks using the 400 microwells/mould and for 2 days using the 1,889 microwells/mould.

### 2.6 Microtissue diameter and sphericity measurements

iMPC and oMPC microtissues were monitored and images were acquired using the 4x objective of an optical inverted microscope (Primo Vert, Zeiss, USA) equipped with a digital camera. Width and length were measured using the ImageJ 1.53e software (Wayne Rasband and Contributors, USA). A diameter ratio of each microtissue was obtained by dividing width by length (sphericity). The analysis was performed using one cell donor and three independent experiments using fifteen spheroids from each group.

### 2.7 Fusion assays

The capacity to form fibrocartilage zonally defined assembled microtissues by spontaneous self-assembly was determined by manually seeding the iMPC, oMPC and MSC microtissues in custom ultrapure non-adherent agarose (Sigma) moulds. The moulds were created using sterile 2% w/v agarose cast into a 12-well plate. The central agarose well of the cylindrical mould was 3 mm in diameter and 1.5 mm in depth, while the central agarose well of the ring mould had an inner diameter of 5 mm, an outer diameter of 8 mm, and a depth of 1.5 mm. The total number of microtissues seeded in the cylindrical-shape mould was 1,700 and, in the ring-shape mould was 3,400. Each well was then topped up with 3 mL of induced medium and cultured in physioxic conditions (37 °C in a humidified atmosphere with 5% CO_2_ and 5% O_2_) for up to 4 weeks.

A modified version of this fusion assay was also developed to allow the assembly of iMPC and oMPC derived microtissues in a single graft. To this end, iMPC and oMPC microtissues were fabricated in a density of 2×10^3^ cells/microwell. After 48h, a total of 3,400 iMPC microtissues were manually assembled in the cylindrical mould and the same amount of oMPC microtissues were manually assembled in the ring mould. After 5 days, iMPC and oMPC assembled microtissues were then assembled together in a 2% (w/v) agarose coated 6-well plate, to avoid cell attachment, and a final volume of 6 mL of induced media was added in each well. Constructs were cultured in physioxic conditions (37°C in a humidified atmosphere with 5% CO_2_ and 5% O_2_) for up to 6 weeks.

### 2.8 Chondroitinase-ABC treatment

On day 14, prior to enzymatic treatment, iMPC and oMPC derived constructs were washed three times in hgDMEM. Following which, they were maintained in an enzymatic solution containing 2 U/mL cABC (Sigma-Aldrich) and 0.05 m acetate (Trizma Base, Sigma-Aldrich) activator in hgDMEM for 4 h in physioxic conditions. After the treatment, the samples were washed again three times with hgDMEM to ensure removal of any residual cABC before the addition of fresh induced medium and the continuation of cultivation in physioxic conditions for the remaining 2 weeks.

### 2.9 Histological evaluation

Briefly, samples were fixed using 4% (w/v) paraformaldehyde (PFA) solution (Sigma) overnight at 4 ◦C. After fixation, samples were dehydrated in a graded series of ethanol solutions (70%–100%), cleared in xylene, and embedded in paraffin wax (all Sigma). Tissue sections (5 μm) were taken at the centre of the defect in the transverse plane using a manual microtome (Leica) and rehydrated before staining. Sections were stained with haematoxylin and eosin (HE) for morphology evaluation, 1% (w/v) alcian blue 8GX in 0.1 M hydrochloric acid (HCL) (AB) to visualize sulphated glycosaminoglycan (sGAG) and counter-stained with 0.1% (w/v) nuclear fast red to determine cellular distribution, 0.1% (w/v) picrosirius red (PSR) for collagen deposition, and 1% (w/v) alizarin red (pH 4.1) to identify calcium deposition (all from Sigma). Stained sections were then imaged using an Aperio ScanScope slide scanner (Leica).

### 2.10 Immunohistochemistry analysis

Immunohistochemistry analyses were performed at the end of the culture period to assess differences in the phenotype between the iMPC and oMPC microtissues and derived constructs. The analyses were performed for collagen I (Abcam ab90395 1:400) and collagen II (Santa Cruz sc52658 1:400) as previously described [38].

### 2.11 Polarized light microscopy (PLM) and collagen alignment quantification

Sections stained with picrosirius red staining were imaged using PLM to visualize collagen fibre orientation. Quantification of mean fiber orientation, fiber dispersion, fibre coherency, and the generation of colour maps was carried out using previously established methods [37,39] utilizing the “directionality” feature in ImageJ *software* as well as the OrientationJ plugin.

### 2.12 Biochemical analysis

After harvested, samples were washed 2 times in phenol-free DMEM (pfDMEM, Sigma-Aldrich, Wicklow, Ireland) media and manually dried. Next, 3.88 U/mL of papain enzyme in 100 mM sodium phosphate buffer/5 mM Na2EDTA/10 mM L-cysteine, pH 6.5 (all from Sigma–Aldrich), was used to digest the samples at 60 ◦C for 18 h using a rotator. The completely digested samples were then vortexed for 1 minute and centrifuged for 5 minutes at 650*g*.

The DNA content was quantified immediately after digestion using the Quant-iT™ PicoGreen ® dsDNA Reagent and Kit (Molecular Probes, Biosciences) following the company recommendation steps. The DNA content of each sample was quantified using the Synergy HT multi-detection microplate reader (BioTek Instruments, Inc) at a wavelength of 480 nm.

The amount of glycosaminoglycans (sGAG) was determined using the dimethylmethylene (DMMB) blue dye-binding assay, with a chondroitin sulphate solution (Blyscan, Biocolor Ltd., Carrickfergus, UK) for the standards. The pH of the DMMB was adjusted to 1.5 using a 12 N HCl solution. The samples were read using the Synergy HT multi-detection microplate reader (BioTek Instruments, Inc) at a wavelength of 530 and 590 nm.

The total collagen content was obtained using a chloramine-T assay. Briefly, samples were initially mixed with 38% HCL (Sigma) and incubated at 110 ◦C for 18 h to allow hydrolysis to occur. Samples were subsequently dried in a fume hood and the sediment reconstituted in ultra-pure water. A concentration of 2.82% (w/v) Chloramine T and 0.05% (w/v) 4-(Dimethylamino) benzaldehyde (both Sigma) were then added in the samples and the hydroxyproline content quantified with a *trans*-4-Hydroxy-L-proline (Fluka analytical) standard using a Synergy HT multi-detection microplate reader at a wavelength of 570 nm (BioTek Instruments, Inc). The total collagen content was estimated by measuring hydroxyproline levels and applying a hydroxyproline-to-collagen ratio of 1:7.69.

### 2.13 Scanning electron microscopy (SEM)

Briefly, samples were initially fixed in 3% (w/v) glutaraldehyde for 2 hours at room temperature and washed twice with 1x PBS. Next, chondroitin sulfate and dermatan sulfate were removed by incubating samples overnight in 0.6 U/mL cABC prepared in a buffer of 0.05 M tris HCl and 0.06M sodium acetate at pH 8.0. Hyaluronic acid was then removed by incubating the samples overnight in 20.4 U/mL hyaluronidase (all from Sigma). Samples were next dehydrated in crescent concentrations of ethanol: 30%, 50%, 70%, 90% and 100% of 10 minutes each and dried using critical point. The dried samples were mounted on SEM pin stubs with carbon adhesive discs and coated with gold/palladium for 60 s at a current of 40 mA using a Cressington 208 h sputter coater. Imaging was carried out in a Zeiss ULTRA plus SEM. The diameter of the collagen fibers was measured using ImageJ software.

### 2.14 Statistical analysis

Statistical analysis was performed using GraphPad Prism software (GraphPad Software, CA, USA). A *two-way* ANOVA was performed followed by Tukey’s multiple comparison post-test to assess the differences in diameter and sphericity of the microtissues with time in culture and the biochemical content of the tissues. A non-paired student t-test was performed to compare the differences in biochemical content of iMPC and oMPC grafts. Numerical and graphical results are presented as mean ± standard deviation. Significance was determined when *p <* 0.05.

## 3. RESULTS

### 3.1 Isolation of meniscal progenitor cells from the inner and outer region of the meniscus

Caprine meniscus progenitor cells (MPCs) were isolated based on their ability to adhere to fibronectin-coated plates within 20 minutes after seeding. Both inner and outer MPCs began to spread by day 5, forming colonies by day 7 that continued to expand with time in culture (Figure 1Ai). No obvious differences in cellular morphology were observed between inner and outer progenitors or the mixed population of fibrochondrocytes (FCs). However, MPCs exhibited a greater colony-forming capacity compared to FCs (Figure 1Aii), with MPCs observed to produce larger diameter colonies compared to FCs (Figure 1Bi). Cumulative population doublings demonstrate the strong proliferative capacity of iMPCs and oMPCs, with a linear increase in cumulative population doublings from passage 1 to passage 4 (Figure 1Bii). Both inner and outer MPCs could generate a calcified matrix following 21 days in osteogenic medium, as indicated by positive alizarin red staining. Lipid droplet formation was also observed in MPCs following 21 days in adipogenic media, though with only a few vacuoles. In contrast, iFCs and oFCs did not show a potential to undergo osteogenic or adipogenic differentiation, as evidenced by the absence of mineral deposition and lipid formation under the relevant culture conditions (Figure 1Cii). The fibrochondrogenic capacity of iMPC and oMPC was assessed in pellet culture using passage 3 cells maintained in chondrogenic medium. Both MPC populations exhibited a fibrochondrogenic phenotype, as demonstrated by the histological analysis where strong staining for sGAG and collagen deposition was observed (Figure 1Ci). Notably, MPCs at a higher passage (passage 4) maintained their fibrochondrogenic potential, as evidenced by the strong staining for collagen and sGAGs (data not shown). Immunohistochemical analysis revealed differences between iMPCs and oMPCs in their fibrocartilage phenotype (Figure 1Ci). Tissues generated by iMPCs stained more intensely for collagen type II, while tissues produced by oMPCs were positive for both collagen types I and II. This staining pattern closely mirrors that observed in the collagen staining of native meniscus controls for each respective region. No significant difference in DNA content was observed between iMPC and oMPC derived pellets (Figure 1D). iMPCs secreted significantly higher levels of sGAG compared to oMPCs, again mimicking the zonal differences observed in the native meniscus tissue (*p<.0484) (Figure 1E). No statistical differences in total collagen content were observed in tissues generated by iMPC and oMPC (Figure 1F).

**Figure 1.**
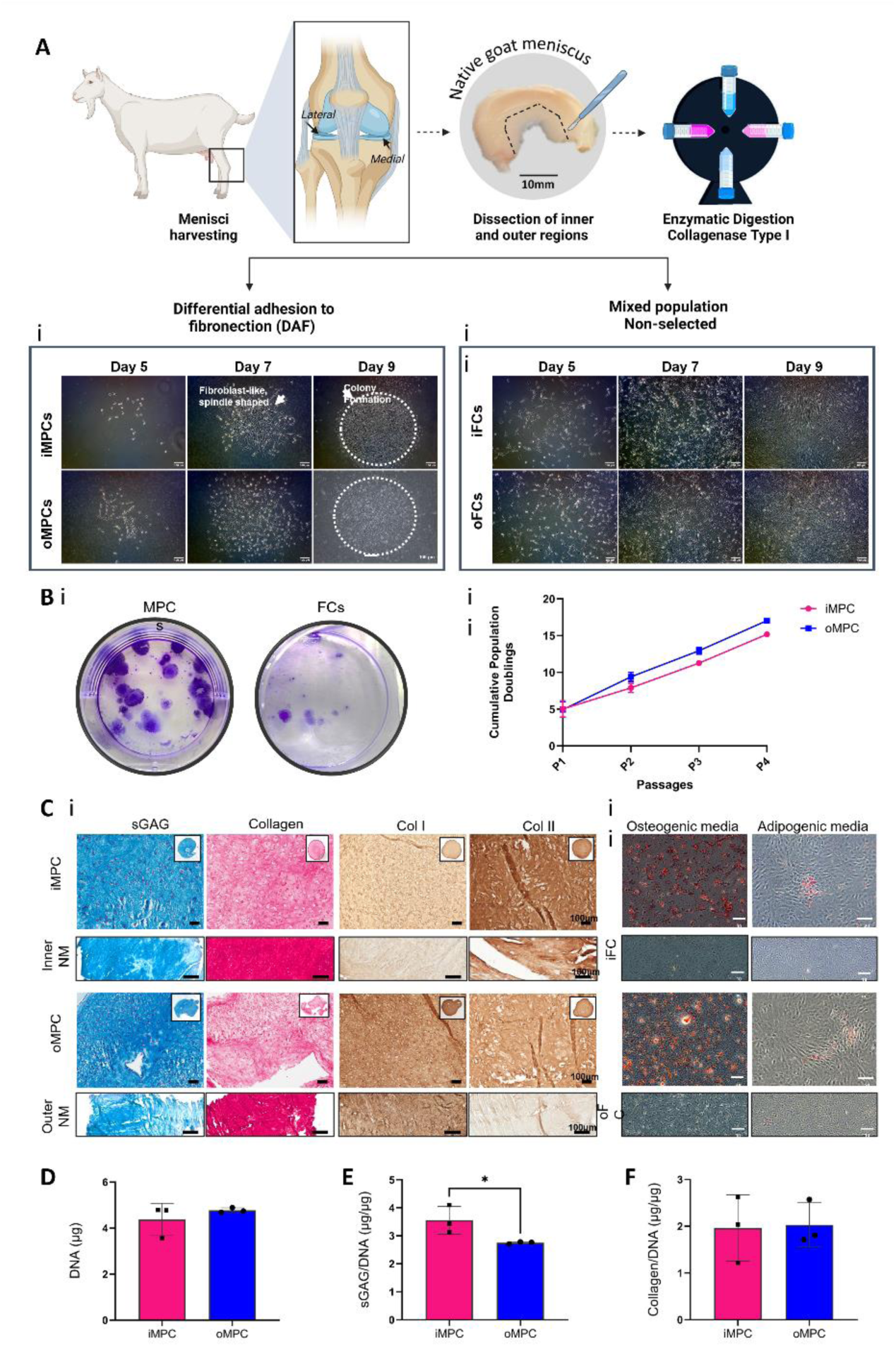
Isolation, morphological comparison, and tri-lineage differentiation potential of inner and outer meniscus progenitor cells (iMPCs and oMPCs). (A) Schematic illustrating the isolation of meniscus progenitor cells (MPCs) and fibrochondrocytes (FCs). Morphological comparison of MPCs (Ai) and FCs (Aii) isolated from goat meniscal tissue at passage 0 (P0) on days 5, 7, and 9 in monolayer culture (scale bar = 100μm). (B) Crystal violet staining to visualise colonies in MPCs and FCs (i). Cumulative population doublings of iMPCs and oMPCs across passages 1, 2, 3, and 4 (ii). (Ci) Histological and immunohistochemical staining of iMPC and oMPC pellets at passage 3: sulphated glycosaminoglycan (Alcian Blue, AB) and total collagens (Picrosirius Red, PR), collagen types I and II after 21 days of chondrogenic culture, with native meniscus (NM) controls (scale bar = 100μm). (Cii) Alizarin red staining for mineralization and Oil Red O staining for lipid deposits in iMPC, oMPC, iFC, and oFCs following 3 weeks of osteogenic and adipogenic differentiation (scale bar = 100μm or 200μm). (D) DNA, (E) sGAG, and (F) collagen content after 21 days of chondrogenic culture. Error bars denote standard deviation. Significance is indicated as follows: *p < .0476, **p < .0028, and ****p < .0001.

### 3.2 Biofabrication of zonally defined MPC derived microtissues

Region-specific meniscal microtissues could potentially be used as biological building blocks for the biofabrication of zonally defined meniscal grafts. To this end, we attempted to generate meniscal microtissues using a range of different starting MPC numbers (1,000, 2,000, and 4,000) per microtissue. On day 0, cells were evenly distributed throughout the microwells, resembling a cell suspension (Figure 2B). By day 2, small cell aggregates were observed in all groups, with microtissue size increasing in proportion to the initial cell number. All groups continued to grow throughout the culture period, with larger aggregates particularly evident for the higher seeding density (4,000 cells per microwell) (Figure 2B). After 21 days of culture, microtissues derived from both iMPCs and oMPCs displayed a uniform and homogeneous morphology at all cell densities, with their size increasing as the initial cell number rose (Figure 2C). Histological staining revealed consistent matrix deposition across groups, with collagen fibers prominently aligned along the edges of the microtissues in a circular pattern. Type II collagen was more pronounced in the iMPC derived microtissues, while type I collagen staining was more intense in oMPC derived microtissues, confirming the development of meniscal microtissues with region-specific characteristics (Figure 2D). sGAG synthesis was significantly higher (p < 0.0151) in the 2000-cell iMPC derived microtissues compared to the 2000 and 4000-cell oMPC derived microtissues (Figure 2F). Total collagen synthesis was not significantly affected by the initial number of MPCs per microtissue or the MPC source (Figure 2G).

**Figure 2:**
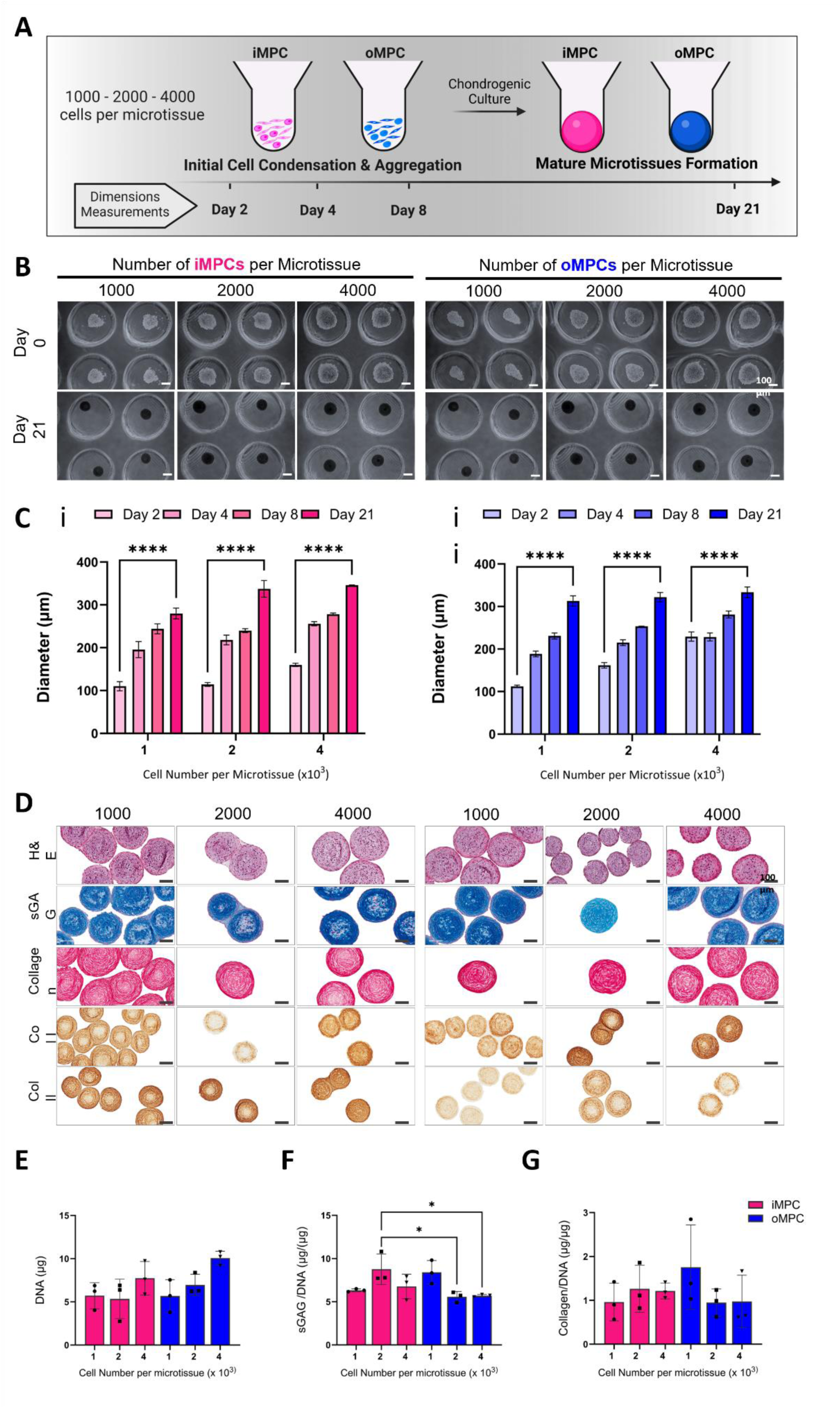
Biofabrication and characterisation of iMPC and oMPC derived microtissues. (A) Schematic illustrating the initial cell seeding densities of iMPC and oMPC in individual microwells for condensation and subsequent microtissue formation using a 401-microwell mould. (B) Microscopic brightfield images of biofabricated iMPC and oMPC derived microtissues at days 4 and 21 during chondrogenic culture, shown at three different cell densities (1000, 2000, and 4000 cell/microtissue) (scale bar = 100μm). (C) Quantification of iMPC (i) and oMPC (ii) derived microtissues diameters over the chondrogenic culture period, error bars denote standard deviation, (n = 3, Mean ± SD), *denotes significance (∗p < .00007). (D) Histological and immunohistochemical analysis of MPCs-derived microtissues after 21 days of chondrogenic induction (scale bar = 100µm). (E) DNA, sGAG, and collagen content after 21 days of chondrogenic differentiation, error bars denote standard deviation, *denotes significance (∗p <.0319).

### 3.3 The fusion of iMPC and oMPC derived microtissues produces phenotypically distinct engineered constructs

The fusion of iMPC and oMPC microtissues in a cylindrical mould was next investigated over a period of 4 weeks in media supplemented with TGF-β3 (Figure 3). iMPC and oMPC derived microtissues were fabricated at densities of 1,000, 2,000 and 3,000 cells/µT. Fusion of both iMPC and oMPC derived microtissues occurred within 24 hours (Figure 3A). Macroscopic images of iMPC and oMPC microtissue derived constructs after 4 weeks of culture show the formation of a compact tissue structure, closely resembling the intended shape defined by the outer mould (Figure 3A).

**Figure 3:**
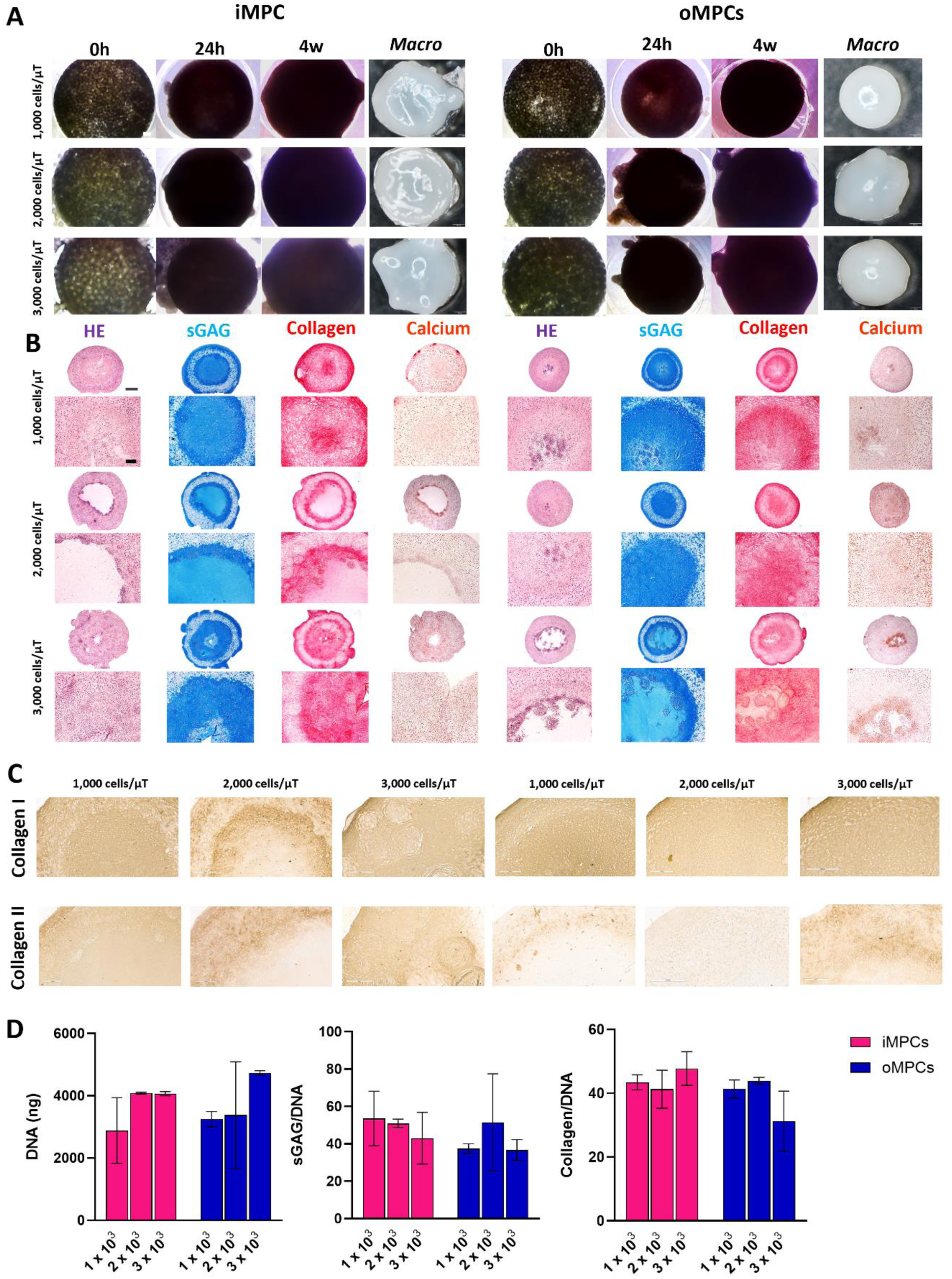
iMPC and oMPC assembled microtissues have differences in phenotype when assembled in the cylindrical shape mould. (A) Phase contrast images of iMPC and oMPC microtissues fabricated using densities of 1×10^3^, 2×10^3^ and 3×10^3^ cells at 0h, 24h and 4 weeks. Macroscopic images of constructs at 4 weeks of culture. (B) Hematoxylin and Eosin (HE), Alcian Blue (sGAG), Picrosirius Red (Collagen) and Alizarin Red (Calcium) stains of iMPC and oMPC assembled microtissues at 4 weeks of culture. (B) Immunohistochemistry of collagens I and II of assembled microtissues at 4 weeks of culture. (C) Biochemical quantification of total DNA, sGAG/DNA and collagen/DNA of iMPC and oMPC assembled microtissues at 4 weeks of culture. Scale bars: (A) – 50 µm; (B) 1 mm and 100 µm; (C) – 200 µm.

The potential of the fused iMPC and oMPC microtissue constructs to form scaled-up fibrocartilage tissue was assessed using histological, immunohistochemical and biochemical analyses (Figure 3B-D). Following 4 weeks of *in vitro* culture, robust fibrochondrogenesis was observed in all iMPC and oMPC derived constructs, as evident by alcian blue staining for sGAG deposition and picrosirius staining for collagen deposition (Figure 3B). Small nodes of calcification were occasionally observed in iMPC and oMPC derived constructs (Figure 3B). Collagen deposition was typically more pronounced in the outer region of the iMPC and oMPC constructs (Figure 3B). The constructs generated using iMPC derived microtissues stained more intensely for collagen type II (Figure 3C). The constructs generated using oMPC microtissues generally stained more homogenously for collagen type I (Figure 3C). Total sGAG and collagen synthesis was comparable in iMPC and oMPC microtissue derived constructs (Figure 3D).

### 3.4 Generation of ring-shaped meniscal grafts using MPC derived microtissues

Generating a large viable tissue with shape fidelity is a key challenge in meniscus tissue engineering. To address this, we seeded iMPC and oMPC derived microtissues into ring-shaped agarose moulds (Figure 4). As a control, we also seeded bone marrow MSC derived microtissues within the same ring mould (supplementary figure 4). iMPC and oMPC microtissues were fabricated at densities of 1,000 and 2,000 cells/µT. After 4 weeks of *in vitro* culture, both iMPC and oMPC derived microtissues fused into a dense ring-shaped tissue (Figure 4A), except for the 1,000 oMPCs per microtissue group which contracted in culture. Histological analysis revealed strong staining for sGAG and no calcium deposition in both iMPC and oMPC ring constructs (Figure 4B). sGAG and collagen deposition, when normalised to DNA content, was significantly higher for constructs generated using 1,000 cells/µT compared to 2,000 cells/µT (Figure 4C). In contrast to that observed using MPCs, MSC derived microtissues fused but contracted into a sphere-like shape by the end of 4-week culture period for both 1,000 and 2,000 cells/µT (supplementary figure 4). The MSC derived microtissues underwent robust chondrogenesis, as indicated by strong staining for sGAG and collagen. It should also be noted that when a lower number of iMPC and oMPC derived microtissues were seeded into the moulds, no ring-shaped tissue formed (supplementary figure 3).

**Figure 4:**
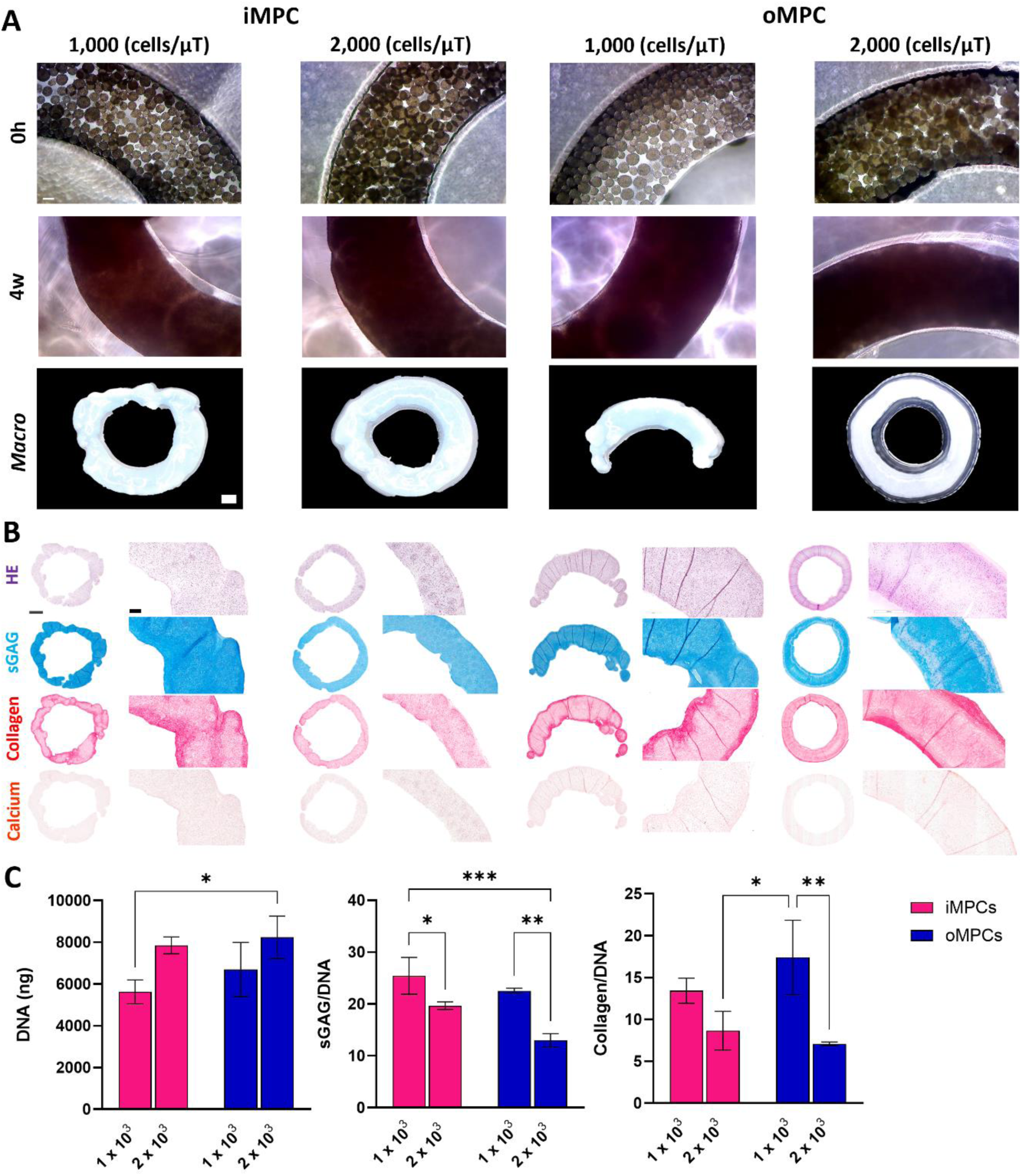
iMPC and oMPC microtissues self-organized in a ring shape and maintain differences in phenotype. (A) Phase contrast images of iMPC and oMPC ring assembled microtissues fabricated using densities of 1×10^3^ and 2×10^3^ cells at 0h and 4 weeks. Macroscopic images of iMPC and oMPC ring assembled microtissues at 4 weeks of culture. (B) Hematoxylin and Eosin (HE), Alcian Blue (sGAG), Picrosirius Red (Collagen) and Alizarin Red (Calcium) stains of iMPC and oMPC ring assembled microtissues at 4 weeks of culture. (C) Immunohistochemistry of collagens I and II of iMPC and oMPC ring assembled microtissues at 4 weeks of culture. (D) Biochemical quantification of total DNA, sGAG/DNA and collagen/DNA of iMPC and oMPC ring assembled microtissues at 4 weeks of culture. The data are expressed as mean ± SD. The asterisks indicate p-values obtained by nonpaired two-way ANOVA followed by Tukey’s multiple comparisons post-test (*p < 0.05; **p < 0.01; ***p < 0.005). Scale bars: (A) – 100 µm and 500 µm (macro); (B) 1 mm and 200 µm.

### 3.5 Enzymatic treatment modulates collagen fiber organization and thickness

Since the use of enzymatic agents has shown great promise in enhancing the quality of engineered cartilage [35,37,60,61] and meniscus [12,14] tissues through modulation of the developing matrix, we next sought to investigate the effect of temporal enzymatic treatment on the functional development of fibrocartilaginous constructs engineered using MPC derived microtissues. To this end, we exposed the tissue rings engineered using MPC derived microtissues to chondroitinase ABC (cABC) for 4 h on day 14 of chondrogenic culture (Figure 5). Enzymatic treatment did not impact the capacity of MPC derived microtissues to generate ring-shaped tissues (Figure 5A). As expected, after 4 weeks of culture, cABC treatment significantly reduced total sGAG levels compared to untreated controls. cABC treatment had no significant effect on DNA or collagen levels (Figure 5C). Untreated and enzymatically treated iMPC and oMPC derived constructs stained positive for type I collagen, while only iMPC derived constructs stained positive for type II collagen (Figure 5D). This data set suggests that the cABC treatment does not affect the phenotype of iMPC and oMPC derived constructs.

**Figure 5:**
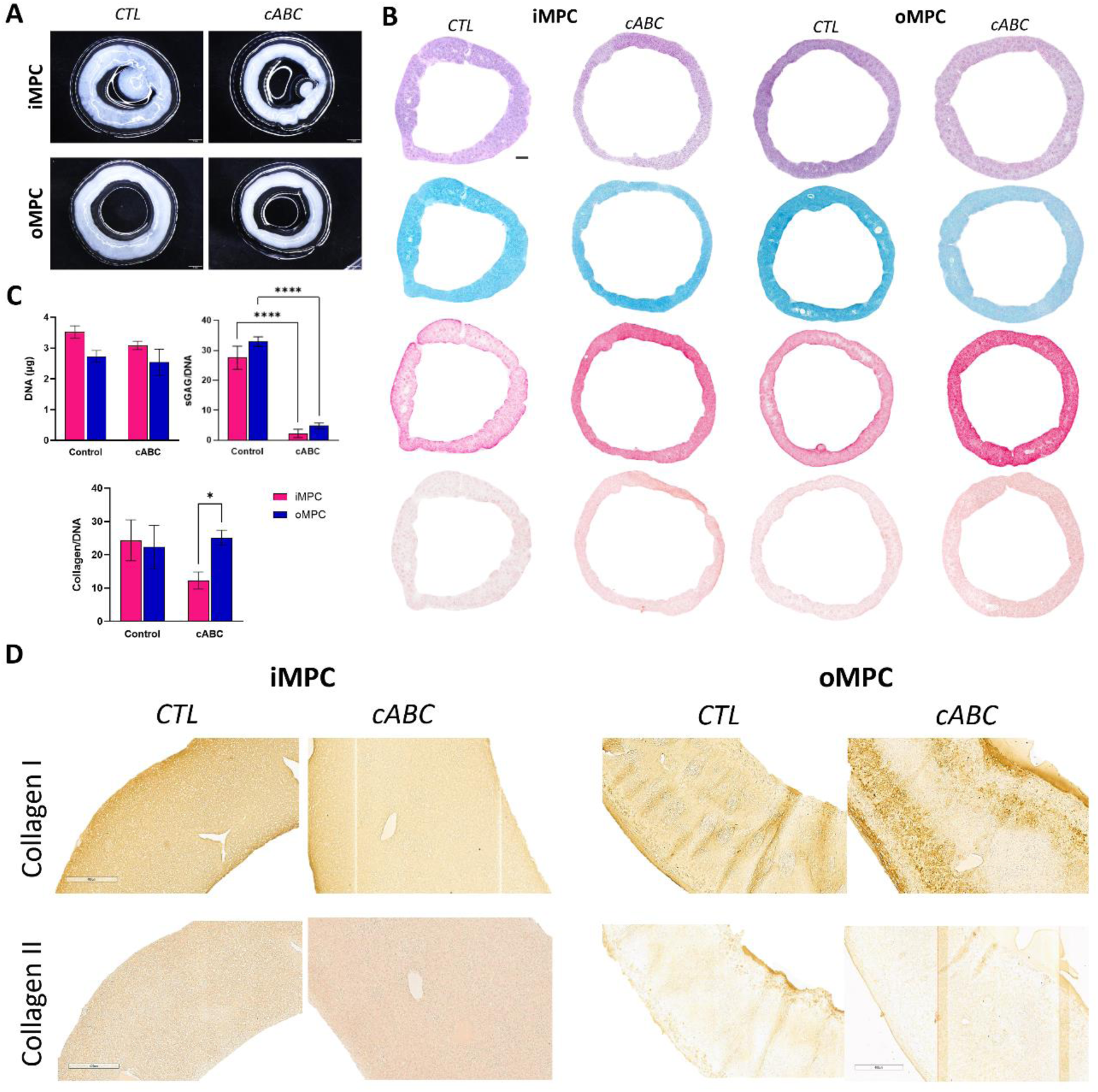
cABC treatment do not impact the phenotype in iMPC and oMPC assembled microtissues constructs. (A) Macroscopic images of untreated and enzymatically treated iMPC and oMPC ring assembled microtissues at 4 weeks of culture. (B) Hematoxylin and Eosin (HE), Alcian Blue (sGAG), Picrosirius Red (Collagen) and Alizarin Red (Calcium) stains of untreated and enzymatically treated iMPC and oMPC ring assembled microtissues at 4 weeks. (C) Biochemical quantification of total DNA, sGAG/DNA and collagen/DNA of untreated and enzymatically treated iMPC and oMPC ring assembled microtissues at 4 weeks. (D) Immunohistochemistry of collagens I and II of untreated and enzymatically treated iMPC and oMPC assembled microtissues at 4 weeks of culture. The data are expressed as mean ± SD. The asterisks indicate p-values obtained by nonpaired two-way ANOVA followed by Tukey’s multiple comparisons post-test (*p < 0.05; **p < 0.01). Scale bars: (A) – 1 mm; (B) – 1 mm and (D) – 400 µm.

Polarized-light microscopy (PLM) was next used to investigate if enzymatic treatment influences collagen organization within the engineered tissue (Figure 6A). Under polarized light microscopy, the colour intensity of collagen fibers showed no noticeable difference between the untreated and cABC treated groups (Figure 6A). However, the coherency values (where a value of 1 indicates fibers are aligned in the same direction, while a value of 0 indicates dispersion of fibers in all directions) were significantly higher for the inner (p < 0.0001) and outer (p < 0.05) assembled microtissues rings treated with the cABC, indicating that the collagen fibers were more aligned (Figure 6C). To further evaluate the influence of cABC treatment on collagen network development and maturation, the iMPC and oMPC assembled microtissues constructs were analysed by SEM (Figure 6B). Enzymatic treatment appeared to enhance collagen alignment compared to untreated controls (Figure 6B). Furthermore, the collagen fiber diameter (Figure 6D) increased from an average of ∼20 nm in untreated samples to an average of ∼100 nm in enzymatically treated groups (p < 0.0001). Collagen fiber diameter was significantly higher in oMPC derived tissues compared to the iMPC derived tissues (p <0.0001).

**Figure 6:**
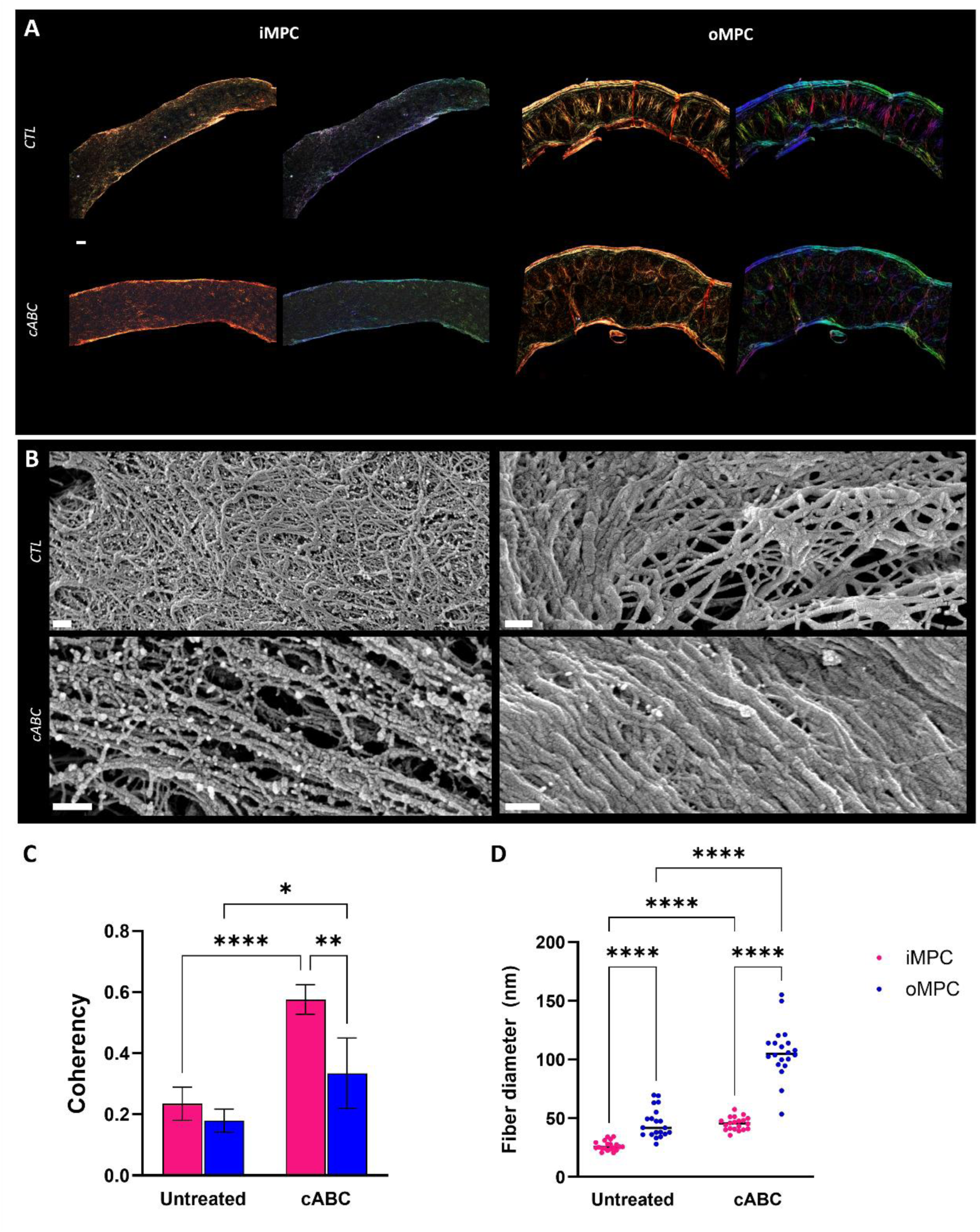
cABC treatment in iMPC and oMPC ring assembled microtissues alters collagen network directionality and increases the thickness of the collagen fibers. (A) Polarized light microscopy (PLM) images of untreated and enzymatically treated iMPC and oMPC ring assembled microtissues at 6 weeks of culture. (B) Scanning electron microscopy (SEM) images of untreated and enzymatically treated iMPC and oMPC ring assembled microtissues at 4 weeks of culture. (C) Collagen fibers coherency quantification, where a value of 1 indicates fibers are aligned in the same direction, while a value of 0 indicates dispersion of fibers in all directions. (D) Quantification of the collagen fiber diameter. The data are expressed as mean ± SD. The asterisks indicate p-values obtained by nonpaired two-way ANOVA followed by Tukey’s multiple comparisons post-test (*p < 0.05; **p < 0.01). Scale bars: (A) – 100 µm; (B) – 200 nm and 300 nm (oMPC under cABC treatment).

### 3.6 The spatial assembly of iMPC and oMPC derived microtissues results in a zonally defined meniscal graft with spatially distinct phenotypes

The meniscus is zonally divided into inner and outer regions with phenotypically different compositions [3], and recapitulation of this zonal organization is believed to be integral to the success of meniscus tissue engineering. Therefore, we next sought to biofabricate a construct with regionally distinct matrix phenotypes resembling the native meniscus by spatially localising iMPC and oMPCs derived microtissues within a supporting mould (Figure 7). To this end we initially separately assembled iMPC and oMPC derived microtissues (2,000 cells/µT), first allowing the iMPC microtissues to fuse in a cylindrically shaped mould, while the oMPC microtissues were allowed to fuse in a ring shape mould. After 5 days of culture, pre-fused iMPC and oMPC microtissue derived constructs were assembled in an agarose coated well and maintained for a further 6 weeks of culture to generate a cohesive graft (graphical abstract figure D).

**Figure 7:**
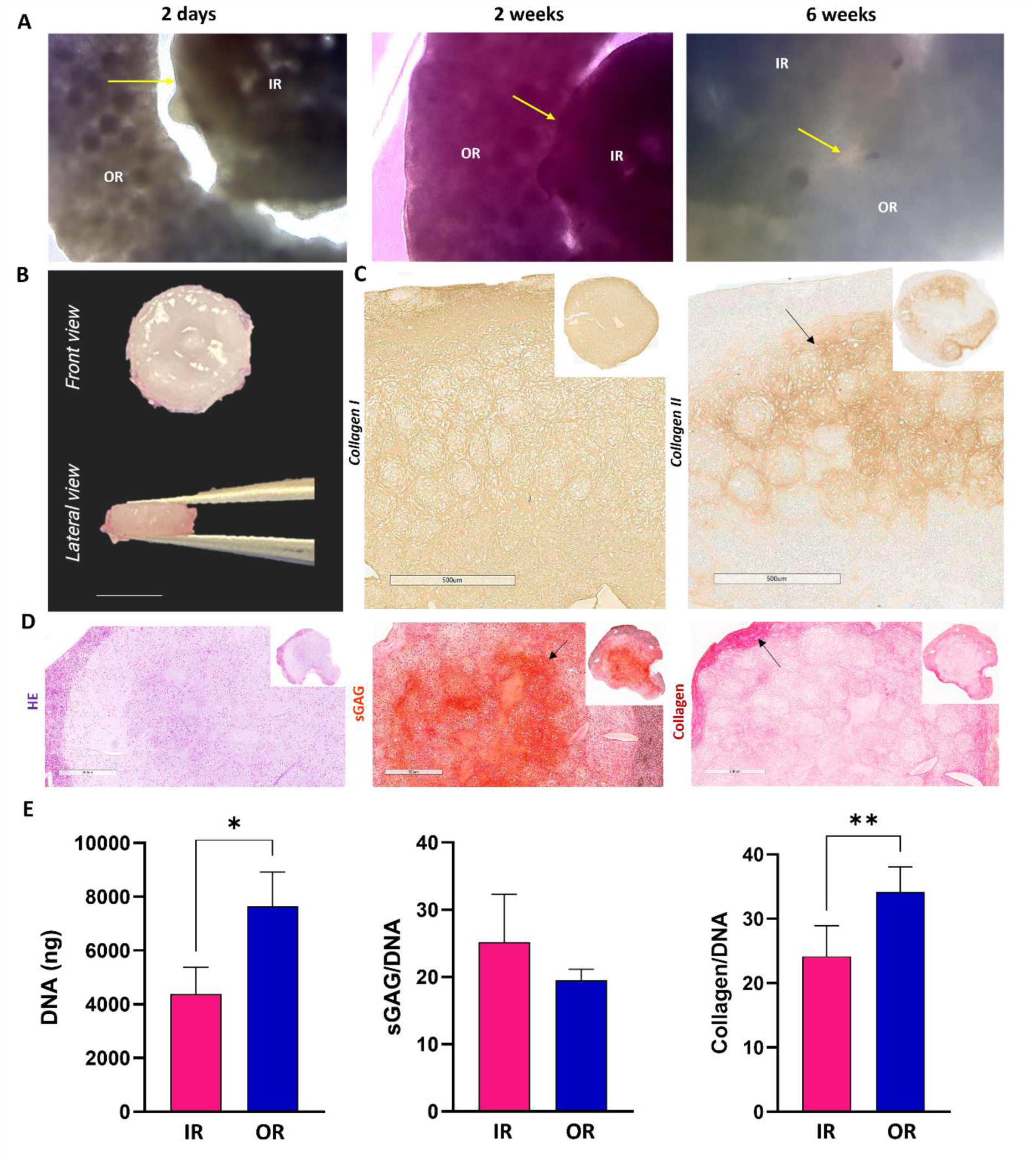
Biofabrication of a zonally defined menisci graft by self-organization of iMPC and oMPC assembled microtissues. (A) Phase contrast images of fused iMPC and oMPC assembled microtissues at 2 days, 2 and 6 weeks. (B) Front and lateral macroscopic images of the zonally defined menisci graft. (C) Immunohistochemistry of collagens I and II of fused iMPC and oMPC assembled microtissues at 6 weeks. Note that the outer region is negative for collagen type II deposition (arrow). (D) Hematoxylin and Eosin (HE), Safranin Red (sGAG) and Picrosirius Red (Collagen) stains of fused iMPC and oMPC assembled microtissues at 6 weeks. Note a stronger intensity of sGAG in the inner core of the construct (arrow) and note a stronger intensity of collagen in the outer region of the construct (arrow). (E) Biochemical quantification of total DNA, sGAG, collagen, sGAG/DNA and collagen/DNA of fused iMPC and oMPC assembled microtissues at 6 weeks. The data are expressed as mean ± SD. The asterisks indicate p-values obtained by nonpaired student t-test (*p < 0.05). Scale bars: (A) – 100 µm; (B) 5 mm; (C and D): 500 µm.

Phase contrast images revealed that there was a noticeable separation between the iMPC and oMPC regions after 2 days of fusion (Figure 7A, arrow), however by the end of the 6 weeks culture period the two zones of the construct were fully integrated (Figure 7A, arrow). Safranin-O staining suggested greater sGAG deposition in the core of the final assembled construct (Figure 7D, arrow) when compared to the outer region. In addition, staining for collagen deposition was more intense in the outer regions of the construct (Figure 7D, arrow). The assembled graft stained positive for type I in both the inner and outer regions (Figure 7C). In contrast, there was no evidence for type II collagen deposition in the outer regions of the assembled graft and in the internal core (Figure 7C, arrow). No significant difference was observed in the sGAG/DNA content; however, the collagen/DNA ratio was significantly higher (p < 0.01) in the oMPC region (Figure 7E).

## 4. DISCUSSION

In this study we report for the first time the biofabrication of zonally defined meniscus grafts through the assembly of phenotypically distinct fibrocartilage microtissues. Microtissues were successfully fabricated using progenitor cells isolated from both the inner (iMPC) and outer (oMPC) regions of the meniscus, with both cell types displaying robust fibrochondrogenesis and distinct meniscus region-specific phenotypes. Furthermore, we observed that iMPC and oMPC derived microtissues could fuse in various spatial configurations, maintaining shape fidelity and generating an extracellular matrix rich in sGAG and collagen type I. We also found that cABC treatment during tissue development modulates collagen fiber formation and organization. Finally, to demonstrate the utility of this biofabrication strategy, we engineered a cohesive yet zonally defined meniscal graft through the assembly of iMPC and oMPC derived assembled microtissues. Such grafts may form the basis of new treatments for damaged or diseased meniscal tissue.

The biofabrication of living meniscal grafts relies on the identification of a suitable cell source with robust fibro-chondrogenic potential. The observation that tissue-resident stem/progenitor cells play a critical role in organ homeostasis and wound healing have motivated their use in tissue engineering and regenerative medicine. Meniscus resident progenitor cells possess an inherent meniscal phenotype and intrinsic regenerative capabilities, motivating their use in emerging meniscus tissue engineering strategies [29]. Similar to previous studies, [14] we found that caprine meniscus progenitor cells can be isolated *via* differential fibronectin adhesion from both the inner (iMPC) and outer (iMPC) regions of the meniscus. These cells exhibit enhanced colony formation, remarkable proliferative capacity and the ability to differentiate towards osteogenic, adipogenic, and (fibro)chondrogenic lineages, while maintaining a zonal fibrochondrogenic phenotype that outperforms that of non-selected fibrochondrocytes (iFC and oFC). This aligns with other studies which highlight the superior clonogenicity, proliferative potential, and multipotent differentiation ability of MPCs from vascular and avascular regions compared to the non-fibronectin adherent meniscus cell population which lack these capabilities [30,40].

Next, we explored the potential of MPCs in the biofabrication of region-specific microtissues with the goal of engineering meniscal grafts capable of mimicking the unique zonal nature of the native tissue. Our study demonstrated that both iMPC and oMPC can be used to biofabricate microtissues that recapitulate certain region-specific characteristics of the native meniscus. We initially explored different cell densities, and as expected, increasing the initial cell numbers led to the formation of larger microtissues. Previous studies also investigated the influence of cell density on hyaline cartilage microtissue properties, highlighting how higher cell densities can enhance tissue growth.[26] Other scaffold-free meniscus tissue engineering strategies have found that lower initial seeding densities results in superior mechanical and biochemical properties, with improved collagen and glycosaminoglycan content [41]. These findings suggest that optimizing cell density is integral to improving the functional properties of the engineered meniscal grafts. iMPC microtissues were rich in type II collagen, while oMPC microtissues contained higher levels of type I collagen. Additionally, higher sGAG synthesis was observed in the iMPC derived microtissues. Such zonal differentiation in collagen expression aligns with previous studies showing that MPCs from inner and outer zones can generate anisotropic fibrocartilaginous tissues with distinct phenotypes when bioprinted within melt electrowritten scaffolds [14].

One of the key advantages of using microtissues for tissue engineering applications is their capability to fuse and generate scaled-up grafts [42,43,44]. After the characterization of iMPC and oMPC microtissues, we evaluated their potential as building-blocks in the bioassembly of larger grafts. When iMPC and oMPC microtissues were assembled into a ring-shaped agarose mold, they generated a tissue with a circumferential aligned collagen network similar to that found in the native meniscus [45,46]. These circumferential collagen fibers resist the tensile forces within the meniscus during knee loading [47]. The periphery of these scaled-up grafts was richer in collagen, which is potentially linked to spatial differences in nutrient availability or biomechanical cues (e.g. matrix tension) that develop within the graft over the culture period. As expected, bone marrow MSC microtissue derived constructs contracted in culture [48,49] and did not fuse into the desired ring shape. Together, these results highlight the benefit of using MPC derived microtissues for engineering meniscal grafts. Additionally, modular assembly strategies such as those described here have high potential for automation, which could help meet the clinical demand for scaled-up grafts [42].

Although microtissue assembly enables the biofabrication of extracellular-matrix rich constructs at a millimeter scale, cohesive fusion between the microtissues resulting in the development of grafts with biomimetic extracellular matrix organization remains a challenge. As previously discussed, the extracellular matrix of the meniscus has an intricate organization that is integral to its function.[50] In this study, we demonstrate that the temporal introduction of an exogenous remodeling enzyme, chondroitinase-ABC (cABC), enhanced the intensity of collagen staining, the diameter of collagen fibrils and the overall organization of the collagen network within the engineered tissue. Numerous studies have explored the use of enzymatic treatments for meniscus engineering [11,12,36,51,52]. In line with our findings, Gonzeles-Leon and collaborators (2020) observed a decrease in sGAG content in the groups treated with cABC, while MacBarb and collaborators (2013) observed the formation of thicker and more organized collagen fibers. Furthermore, Huey and collaborators (2011) observed an improvement in collagen organization with cABC treatment. Other approaches that could be adopted in the future include the addition of an exogenous collagen crosslinking agents such as lysyl oxidase (LOX-2), in conjunction with cABC, to improve collagen fibril maturity and tissue functionality [12] Taken together, our data suggest a potential benefit in applying cABC treatment to meniscus grafts engineered using MPC derived microtissues. Future studies will explore the influence of cABC treatment on the biomechanical properties of such grafts.

Despite numerous studies aimed at developing constructs for meniscus repair, [53,54,55,56] replicating the meniscus’ heterogeneous composition, including its zonal anatomy, cell phenotype, and region-specific composition in a single graft remains challenging [57]. Previously, we engineered zonally defined meniscus constructs by inkjeting porcine iMPC and oMPC suspensions into MEW scaffolds, resulting in a graft with native-like extracellular matrix composition and promising mechanical properties [14]. In this study, we report the novel assembly of iMPC and oMPC derived microtissues to create an integrated meniscus graft. The final construct displayed heterogeneous extracellular matrix composition and biochemical content, mimicking some aspects of the native meniscus. Previous studies have demonstrated the potential of modular assembly techniques for engineering tissues like cartilage [24], bone [58], and vessels [59], highlighting the potential of microtissues as building blocks for hierarchical tissue engineering. Our results further support this approach, demonstrating that the spatial assembly of iMPC and oMPC derived microtissues supports the development a meniscus-like graft, paving the way for preclinical studies on meniscus repair.

In conclusion, this study demonstrates the successful biofabrication of zonally defined meniscus grafts using MPC derived meniscal microtissues. iMPCs and oMPCs isolated from the inner and outer meniscus were highly proliferative and displayed distinct phenotypes. Both iMPC and oMPC derived microtissues were able to fuse and develop region-specific meniscus tissue when cast into cylindrical and ring-shaped moulds. Enzymatic treatment with cABC enhanced collagen fiber network maturation without altering the phenotype of the constructs, highlighting its potential for optimizing graft properties. The integration of iMPC and oMPC derived microtissues enabled the biofabrication of a scaled-up, zonally defined and cohesive meniscus graft. Overall, these findings highlight the translational potential of meniscal microtissues-derived grafts for meniscus repair applications.

## CONFLICTS OF INTEREST

The authors declare no conflict of interest.

## ACKNOWLEDGMENTS

We would like to thank Dr Megan Canavan from Trinity College Dublin for the SEM imaging. Schematic diagrams of graphical abstract (created by F.D.S), figures 1 and 7 were created with BioRender.com and Autodesk Fusion 360 software.

## FUNDING

This work was founded by the European Research Council (ERC, 4D-BOUNDARIES #101019344) and Research Ireland Postdoctoral Fellowship (GOIPD/2024/18).

## Notes

### Competing Interest Statement

The authors have declared no competing interest.

## REFERENCES

[1] Fox, A. J., Wanivenhaus, F., Burge, A. J., Warren, R. F., & Rodeo, S. A., The human meniscus: a review of anatomy, function, injury, and advances in treatment, 2015, Clin Anat, 28 (2), 269–287, 10.1002/ca.22456.

[2] Furumatsu, T., Kanazawa, T., Yokoyama, Y., Abe, N., and Ozaki, T. Inner Meniscus Cells Maintain Higher Chondrogenic Phenotype Compared with Outer Meniscus Cells. Connective Tissue Research 52, no. 6 (2011): 459–465. 10.3109/03008207.2011.562061.

[3] Makris, E. A., Hadidi, P., & Athanasiou, K. A., The knee meniscus: structure-function, pathophysiology, current repair techniques, and prospects for regeneration, 2011, Biomaterials, 32(30), 7411–7431, 10.1016/j.biomaterials.2011.06.037.

[4] Twomey-Kozak, J., & Jayasuriya, C. T., Meniscus repair and regeneration: A systematic review from a basic and translational science perspective, 2020, Clin Sports Med, 39(1), 125–163, 10.1016/j.csm.2019.08.003.

[5] Ding, G., Du, J., Hu, X., & Ao, Y., Mesenchymal stem cells from different sources in meniscus repair and regeneration, 2022, Front Bioeng Biotechnol, 10, 796367, 10.3389/fbioe.2022.796367.

[6] Ozeki, N., Koga, H., and Sekiya, I. Degenerative Meniscus in Knee Osteoarthritis: From Pathology to Treatment. Life 12, no. 4 (2022): 603. 10.3390/life12040603.

[7] Baynat, C., Andro, C., Vincent, J.P., Schiele, P., Buisson, P., Dubrana, F., and Gunepin, F.X. Actifit® Synthetic Meniscal Substitute: Experience with 18 Patients in Brest, France. Orthopaedics & Traumatology: Surgery & Research 100, no. 8, Suppl. (2014): S385–S389. 10.1016/j.otsr.2014.09.007.

[8] Merkely, G., Minas, T., Ogura, T., Ackermann, J., Barbieri Mestriner, A., and Gomoll, A. H. Safety, feasibility, and radiographic outcomes of the anterior meniscal takedown technique to approach chondral defects on the tibia and posterior femoral condyle: a matched control study. Cartilage 12, no. 1 (2021): 62–69. 10.1177/1947603518809409.

[9] Ranmuthu, C. D. S., Ranmuthu, C. K. I., Russell, J. C., Singhania, D., and Khan, W. S. Are the Biological and Biomechanical Properties of Meniscal Scaffolds Reflected in Clinical Practice? A Systematic Review of the Literature. International Journal of Molecular Sciences 20, no. 3 (2019): 632. 10.3390/ijms20030632.

[10] Tarafder, S., Ghataure, J., Langford, D., Brooke, R., Kim, R., Eyen, S. L., Bensadoun, J., Felix, J. T., Cook, J. L., and Lee, C. H. Advanced Bioactive Glue Tethering Lubricin/PRG4 to Promote Integrated Healing of Avascular Meniscus Tears. Bioactive Materials 28 (2023): 61–73. 10.1016/j.bioactmat.2023.04.026.

[11] Gonzalez-Leon, E. A., Bielajew, B. J., Hu, J. C., and Athanasiou, K. A. Engineering Self-Assembled Neomenisci through Combination of Matrix Augmentation and Directional Remodeling. Acta Biomaterialia 109 (2020): 73–81. 10.1016/j.actbio.2020.04.019.

[12] Bates, M. E., Troop, L., Brown, M. E., and Puetzer, J. L. Temporal Application of Lysyl Oxidase During Hierarchical Collagen Fiber Formation Differentially Affects Tissue Mechanics. Acta Biomaterialia 160 (2023): 98–111. 10.1016/j.actbio.2023.02.024.

[13] Bernal, P. N., Delrot, P., Loterie, D., Li, Y., Malda, J., Moser, C., and Levato, R. Volumetric Bioprinting of Complex Living-Tissue Constructs within Seconds. Advanced Materials 31, no. 42 (2019): e1904209. 10.1002/adma.201904209.

[14] Barceló, X., Eichholz, K., Gonçalves, I., Kronemberger, G. S., Dufour, A., Garcia, O., Kelly, D. J. Bioprinting of Scaled-Up Meniscal Grafts by Spatially Patterning Phenotypically Distinct Meniscus Progenitor Cells within Melt Electrowritten Scaffolds. Biofabrication 16 (2024): 015013.

[15] Wang, B., Barceló, X., Von Euw, S., and Kelly, D. J. 3D Printing of Mechanically Functional Meniscal Tissue Equivalents Using High Concentration Extracellular Matrix Inks. Materials Today Bio 20 (2023): 100624. 10.1016/j.mtbio.2023.100624.

[16] Zhang, Z. Z., Chen, Y. R., Wang, S. J., Zhao, F., Wang, X. G., Yang, F., Shi, J. J., Ge, Z. G., Ding, W. Y., Yang, Y. C., Zou, T. Q., Zhang, J. Y., Yu, J. K., and Jiang, D. Orchestrated Biomechanical, Structural, and Biochemical Stimuli for Engineering Anisotropic Meniscus. Science Translational Medicine 11, no. 487 (2019): eaao0750. 10.1126/scitranslmed.aao0750.

[17] Dufour, A., Essayan, L., Thomann, C., Petiot, E., Gay, I., Barbaroux, M., and Marquette, C. Confined Bioprinting and Culture in Inflatable Bioreactor for the Sterile Bioproduction of Tissues and Organs. Scientific Reports 14, no. 1 (2024): 11003. 10.1038/s41598-024-60382-2.

[18] Nichol, J.W., and Khademhosseini, A. Modular Tissue Engineering: Engineering Biological Tissues from the Bottom Up. Soft Matter 5, no. 7 (2009): 1312–1319. 10.1039/b814285h.

[19] Ouyang, L., Armstrong, J. P. K., Salmeron-Sanchez, M., and Stevens, M. M. Assembling Living Building Blocks to Engineer Complex Tissues. Advanced Functional Materials (2020). 10.1002/adfm.201909009.

[20] Kronemberger GS, Dalmônico GML, Rossi AL, Leite PEC, Saraiva AM, Beatrici A, Silva KR, Granjeiro JM, Baptista LS. Scaffold- and serum-free hypertrophic cartilage tissue engineering as an alternative approach for bone repair. Artif Organs. 2020 Jul;44(7):E288–E299. 10.1111/aor.13637.

[21] Kronemberger GS, Beatrici A, Dalmônico GML, Rossi AL, Miranda GASC, Boldrini LC, Mauro Granjeiro J, Baptista LS. The hypertrophic cartilage induction influences the building-block capacity of human adipose stem/stromal cell spheroids for biofabrication. Artif Organs. 2021 Oct;45(10):1208–1218. 10.1111/aor.14000.

[22] Nulty J, Freeman FE, Browe DC, Burdis R, Ahern DP, Pitacco P, Lee YB, Alsberg E, Kelly DJ. 3D bioprinting of prevascularised implants for the repair of critically-sized bone defects. Acta Biomater. 2021 May;126:154–169. 10.1016/j.actbio.2021.03.003.

[23] Schott, N. G., Vu, H., and Stegemann, J. P. Multimodular Vascularized Bone Construct Comprised of Vasculogenic and Osteogenic Microtissues. Biotechnology and Bioengineering 119, no. 11 (2022): 3284–3296. 10.1002/bit.28201.

[24] De Moor L, Minne M, Tytgat L, Vercruysse C, Dubruel P, Van Vlierberghe S, Declercq H. Tuning the Phenotype of Cartilage Tissue Mimics by Varying Spheroid Maturation and Methacrylamide-Modified Gelatin Hydrogel Characteristics. Macromol Biosci. 2021 May;21(5):e2000401. 10.1002/mabi.202000401.

[25] Burdis, R., Chariyev-Prinz, F., and Kelly, D. J. Bioprinting of Biomimetic Self-Organised Cartilage with a Supporting Joint Fixation Device. Biofabrication 14, no. 1 (November 25, 2021). 10.1088/1758-5090/ac36be.

[26] Burdis, R., Chariyev-Prinz, F., Browe, D. C., Freeman, F. E., Nulty, J., McDonnell, E. E., Eichholz, K. F., Wang, B., Brama, P. and Kelly, D. J. Spatial Patterning of Phenotypically Distinct Microtissues to Engineer Osteochondral Grafts for Biological Joint Resurfacing. Biomaterials 289 (October 2022): 121750. 10.1016/j.biomaterials.2022.121750.

[27] Kronemberger, G. S., Spagnuolo, F. D., Karam, A. S., Chattahy, K., Storey, K. J., Kelly, D. J. Growth Factor Stimulation Regimes to Support the Development and Fusion of Cartilage Microtissues. Tissue Engineering Part C: Methods 31, no. 1 (January 2025): 36–48. 10.1089/ten.tec.2024.0309.

[28] Spagnuolo, F. D., Kronemberger, G. S., Storey, K. J., and Kelly, D. J. The Maturation State and Density of Human Cartilage Microtissues Influence Their Fusion and Development into Scaled-Up Grafts. Acta Biomaterialia 194 (March 1, 2025): 109–121. 10.1016/j.actbio.2025.01.024.

[29] Seol, D., Zhou, C., Brouillette, M.J., Song, I., Yu, Y., Choe, H.H., Lehman, A.D., Jang, K.W., Fredericks, D.C., Laughlin, B.J., and Martin, J.A. Characteristics of Meniscus Progenitor Cells Migrated from Injured Meniscus. Journal of Orthopaedic Research 35, no. 9 (2017): 1966–1972. 10.1002/jor.23472.

[30] Wang, J., Roberts, S., Li, W., and Wright, K. Phenotypic Characterization of Regional Human Meniscus Progenitor Cells. Frontiers in Bioengineering and Biotechnology 10 (2022): 1003966. 10.3389/fbioe.2022.1003966.

[31] Yan, W.T., Wang, J.S., Guo, S.Y., Zhu, J.H., and Zhang, Z.Z. Isolation and Characterization of Meniscus Progenitor Cells From Rat, Rabbit, Goat, and Human. Cartilage (2024): 19476035241266579. 10.1177/19476035241266579.

[32] Makris, E. A., et al. Combined Use of Chondroitinase-ABC, TGF-β1, and Collagen Crosslinking Agent Lysyl Oxidase to Engineer Functional Neotissues for Fibrocartilage Repair. Biomaterials 35, no. 25 (2014): 6787–6796.

[33] Kwon, H., et al. Translating the Application of Transforming Growth Factor-β1, Chondroitinase-ABC, and Lysyl Oxidase-Like 2 for Mechanically Robust Tissue-Engineered Human Neocartilage. Journal of Tissue Engineering and Regenerative Medicine 13, no. 2 (2019): 283–294.

[34] O’Connell, G.D., Nims, R.J., Green, J., Cigan, A.D., Ateshian, G.A., and Hung, C.T. Time and Dose-Dependent Effects of Chondroitinase ABC on Growth of Engineered Cartilage. European Cells and Materials 27 (2014): 312–320. 10.22203/ecm.v027a22.

[35] Bautista, C.A., H.J. Park, C.M. Mazur, R.K. Aaron, and B. Bilgen. Effects of Chondroitinase ABC-Mediated Proteoglycan Digestion on Decellularization and Recellularization of Articular Cartilage. PLOS One 11, no. 7 (2016): e0158976. 10.1371/journal.pone.0158976.

[36] Lopez, S. G., Kim, J., Estroff, L. A., and Bonassar, L. J. Removal of GAGs Regulates Mechanical Properties, Collagen Fiber Formation, and Alignment in Tissue Engineered Meniscus. ACS Biomaterials Science & Engineering 9, no. 3 (2023): 1608–1619. 10.1021/acsbiomaterials.3c00136.

[37] Burdis R, Gallostra XB, Kelly DJ. Temporal Enzymatic Treatment to Enhance the Remodeling of Multiple Cartilage Microtissues into a Structurally Organized Tissue. Adv Healthc Mater. 2024 Jan;13(3):e2300174. 10.1002/adhm.202300174.

[38] Buckley CT, Vinardell T, Kelly DJ. Oxygen tension differentially regulates the functional properties of cartilaginous tissues engineered from infrapatellar fat pad derived MSCs and articular chondrocytes. Osteoarthritis Cartilage. 2010 Oct;18(10):1345–54. doi: 10.1016/j.joca.2010.07.004. Epub 2010 Aug 3. PMID: 20650328.

[39] Rezakhaniha, R., Agianniotis, A., Schrauwen, J.T., Griffa, A., Sage, D., Bouten, C.V., van de Vosse, F.N., Unser, M., and Stergiopulos, N. Experimental Investigation of Collagen Waviness and Orientation in the Arterial Adventitia Using Confocal Laser Scanning Microscopy. Biomechanics and Modeling in Mechanobiology 11, no. 3-4 (2012): 461–473. 10.1007/s10237-011-0325-z.

[40] Pattappa, G., F. Reischl, J. Jahns, R. Schewior, S. Lang, J. Zellner, B. Johnstone, D. Docheva, and P. Angele. Fibronectin adherent cell populations derived from avascular and vascular regions of the meniscus have enhanced clonogenicity and differentiation potential under physioxia. Frontiers in Bioengineering and Biotechnology 9 (2022): 789621. 10.3389/fbioe.2021.789621.

[41] Hadidi, P., Yeh, T. C., Hu, J. C. H., Athanasiou, K. A. Critical Seeding Density Improves the Properties and Translatability of Self-Assembling Anatomically Shaped Knee Menisci. Acta Biomaterialia 11 (2015): 41–51. doi: 10.1016/j.actbio.2014.09.011.

[42] Schon, B. S., Hooper, G. J., and Woodfield, T. B. Modular Tissue Assembly Strategies for Biofabrication of Engineered Cartilage. Annals of Biomedical Engineering 45, no. 1 (2017): 100–114. 10.1007/s10439-016-1609-3.

[43] Parfenov, V. A., Koudan, E. V., Krokhmal, A. A., et al. Biofabrication of a Functional Tubular Construct from Tissue Spheroids Using Magnetoacoustic Levitational Directed Assembly. Advanced Healthcare Materials 9, no. 24 (2020): e2000721. 10.1002/adhm.202000721.

[44] Schneider, O., Moruzzi, A., Fuchs, S., Grobel, A., Schulze, H. S., Mayr, T., and Loskill, P. Fusing Spheroids to Aligned μ-Tissues in a Heart-on-Chip Featuring Oxygen Sensing and Electrical Pacing Capabilities. Materials Today Bio 15 (2022): 100280. 10.1016/j.mtbio.2022.100280.

[45] Fisher, M. B., Henning, E. A., Söegaard, N., Bostrom, M., Esterhai, J. L., and Mauck, R. L. Engineering Meniscus Structure and Function via Multi-Layered Mesenchymal Stem Cell-Seeded Nanofibrous Scaffolds. Journal of Biomechanics 48, no. 8 (2015): 1412–1419. 10.1016/j.jbiomech.2015.02.036.

[46] Patel, J. M., Brzezinski, A., Ghodbane, S. A., Tarapore, R., Lu, T. M., Gatt, C. J., and Dunn, M. G. Personalized Fiber-Reinforcement Networks for Meniscus Reconstruction. Journal of Biomechanical Engineering 142, no. 5 (2020): 051008. 10.1115/1.4045402.

[47] Bansal, S., Mandalapu, S., Aeppli, C., Qu, F., Szczesny, S. E., Mauck, R. L., and Zgonis, M. H. “Mechanical Function Near Defects in an Aligned Nanofiber Composite Is Preserved by Inclusion of Disorganized Layers: Insight into Meniscus Structure and Function. Acta Biomaterialia 56 (2017): 102–109. 10.1016/j.actbio.2017.01.074.

[48] Vickers, S. M., Gotterbarm, T., and Spector, M. Cross-Linking Affects Cellular Condensation and Chondrogenesis in Type II Collagen-GAG Scaffolds Seeded with Bone Marrow-Derived Mesenchymal Stem Cells. Journal of Orthopaedic Research 28, no. 9 (2010): 1184–1192. 10.1002/jor.21113.

[49] Hilton, S. A., Dewberry, L. C., Hodges, M. M., Hu, J., Xu, J., Liechty, K. W., and Zgheib, C. Mesenchymal Stromal Cells Contract Collagen More Efficiently Than Dermal Fibroblasts: Implications for Cytotherapy. PLOS One 14, no. 7 (2019): e0218536. 10.1371/journal.pone.0218536.

[50] Li, Q., Qu, F., Han, B., Wang, C., Li, H., Mauck, R.L., and Han, L. Micromechanical Anisotropy and Heterogeneity of the Meniscus Extracellular Matrix. Acta Biomaterialia 54 (2017): 356–366. 10.1016/j.actbio.2017.02.043.

[51] Huey, D. J., and Athanasiou, K. A. Maturational Growth of Self-Assembled, Functional Menisci as a Result of TGF-β1 and Enzymatic Chondroitinase-ABC Stimulation. Biomaterials 32, no. 8 (2011): 2052–2058. 10.1016/j.biomaterials.2010.11.041.

[52] MacBarb, R. F., Makris, E. A., Hu, J. C., and Athanasiou, K. A. A Chondroitinase-ABC and TGF-β1 Treatment Regimen for Enhancing the Mechanical Properties of Tissue-Engineered Fibrocartilage. Acta Biomaterialia 9, no. 1 (2013): 4626–4634. 10.1016/j.actbio.2012.09.037.

[53] Zhang, Z.Z., Jiang, D., Ding, J.X., Wang, S.J., Zhang, L., Zhang, J.Y., Qi, Y.S., Chen, X.S., and Yu, J.K. Role of Scaffold Mean Pore Size in Meniscus Regeneration. Acta Biomaterialia 43 (2016): 314–326. 10.1016/j.actbio.2016.07.050.

[54] Yang, Y., Chen, Z., Song, X., Zhang, Z., Zhang, J., Shung, K. K., Zhou, Q., and Chen, Y. Biomimetic Anisotropic Reinforcement Architectures by Electrically Assisted Nanocomposite 3D Printing. Advanced Materials 29 (2017): 1605750. 10.1002/adma.201770076.

[55] Bahcecioglu, G., Bilgen, B., Hasirci, N., et al. Anatomical Meniscus Construct with Zone-Specific Biochemical Composition and Structural Organization. Biomaterials 218 (2019): 119361.

[56] Xu, B., Ye, J., Fan, B.S., Wang, X., Zhang, J.Y., Song, S., Song, Y., Jiang, W.B., Wang, X., and Yu, J.K. Protein-Spatiotemporal Partition Releasing Gradient Porous Scaffolds and Anti-Inflammatory and Antioxidant Regulation Remodel Tissue Engineered Anisotropic Meniscus. Bioactive Materials 20 (2022): 194–207. 10.1016/j.bioactmat.2022.05.019.

[57] Du, M.Z., Dou, Y., Ai, L.Y., Su, T., Zhang, Z., Chen, Y.R., and Jiang, D. Meniscus Heterogeneity and 3D-Printed Strategies for Engineering Anisotropic Meniscus. International Journal of Bioprinting 9, no. 3 (2023): 693. 10.18063/ijb.693.

[58] Hall, G.N., Chandrakar, A., Pastore, A., Ioannidis, K., Moisley, K., Cirstea, M., Geris, L., Moroni, L., Luyten, F.P., Wieringa, P., and Papantoniou, I. Engineering Bone-Forming Biohybrid Sheets through the Integration of Melt Electrowritten Membranes and Cartilaginous Microspheroids. Acta Biomaterialia 165 (2023): 111–124. 10.1016/j.actbio.2022.10.037.

[59] Itoh, M., Nakayama, K., Noguchi, R., Kamohara, K., Furukawa, K., Uchihashi, K., et al. Scaffold-Free Tubular Tissues Created by a Bio-3D Printer Undergo Remodeling and Endothelialization When Implanted in Rat Aortae. PLOS One 10 (2015): e0136681. 10.1371/journal.pone.0136681.

[60] Natoli, R.M., Revell, C.M., and Athanasiou, K.A. Chondroitinase ABC Treatment Results in Greater Tensile Properties of Self-Assembled Tissue-Engineered Articular Cartilage. Tissue Engineering Part A 15, no. 10 (2009): 3119–3128. 10.1089/ten.TEA.2008.0478.

[61] Link, J.M., Hu, J.C., and Athanasiou, K.A. Chondroitinase ABC Enhances Integration of Self-Assembled Articular Cartilage, but Its Dosage Needs to Be Moderated Based on Neocartilage Maturity. Cartilage 13, no. 2_suppl (2021): 672S–683S. 10.1177/1947603520918653.

