## Supplementary figures 1-6 for "Bioassembly of region-specific fibrocartilage microtissues to engineer zonally defined meniscal grafts"

**Key-words**: microtissues, building-blocks, progenitor cell, meniscus, bioassembly, biofabrication.

#Corresponding Author

Daniel J. Kelly − Trinity Centre for Biomedical Engineering, Trinity Biomedical Sciences Institute, Trinity College Dublin, Dublin D02 R590, Ireland; Department of Mechanical, Manufacturing and Biomedical Engineering, School of Engineering, Trinity College Dublin, Dublin D02 R590, Ireland; Department of Anatomy and Regenerative Medicine, Royal College of Surgeons in Ireland, Dublin D02 YN77, Ireland; Advanced Materials and Bioengineering Research Centre (AMBER), Royal College of Surgeons in Ireland and Trinity College Dublin, Dublin D02 F6N2, Ireland; orcid.org/0000-0003-4091-0992; Phone: +353-1-8963947;.

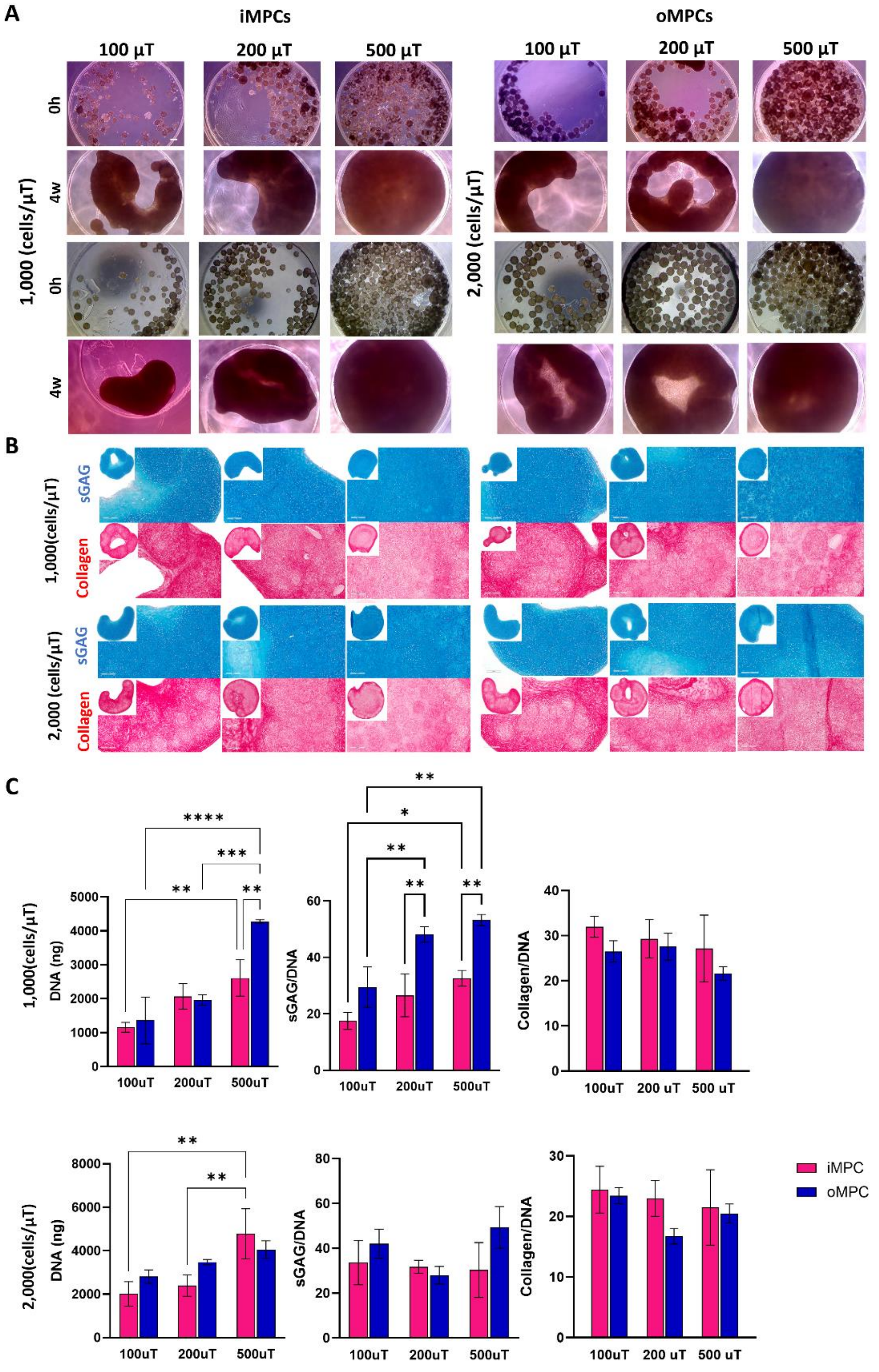

***Supplementary figure 1: iMPC and oMPC microtissues maintain fuse properties at lower densities and demonstrate higher expansion throughout the culture period.*** *(A) Phase contrast images of approximately 100, 200 and 500 fused iMPC and oMPC microtissues fabricated at the densities of 1x10³ and 2x10e³ cells at 0h and 4 weeks. (B) Alcian Blue (sGAG) and Picrosirius Red (Collagen) stains of iMPC and oMPC assembled microtissues. (C) Biochemical quantification of total DNA, sGAG/DNA and collagen/DNA of iMPC and oMPC assembled microtissues. The data are expressed as mean ± SD. The asterisks indicate p-values obtained by unpaired two-way ANOVA followed by Tukey's multiple comparisons post-test (*p < 0.05; **p < 0.01; ***p < 0.005; ****p < 0.001). Scale bars: (A) – 50 µm and (B) – 200 µm.*

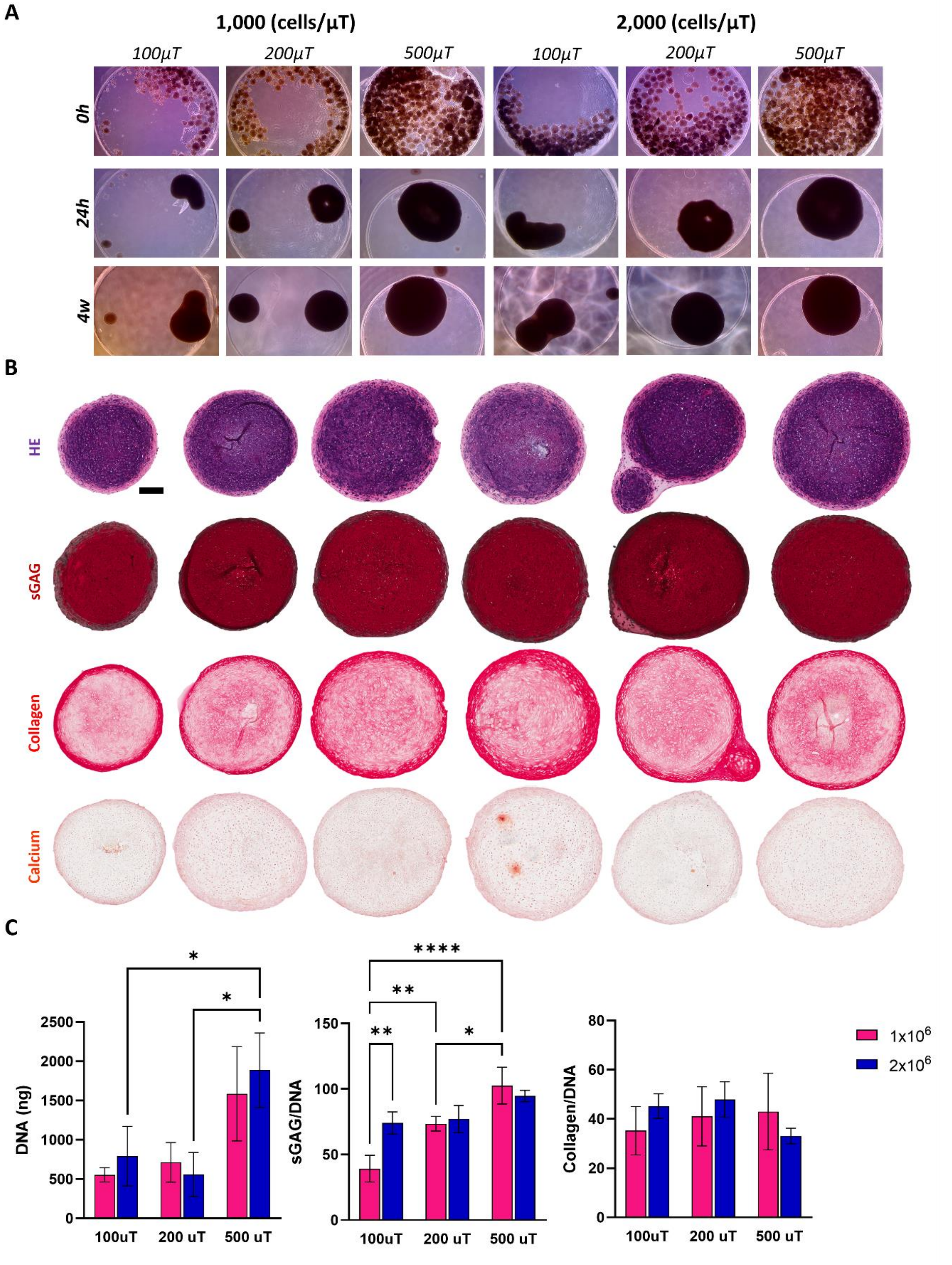


***Supplementary figure 2: MSC microtissues fuse at lower densities and show high contraction.*** *(A) Phase contrast images of approximately 100, 200 and 500 fused MSC microtissues fabricated at the densities of $1x10^3$ and $2x10e^3$ cells at 0h, 24h and 4 weeks. (B) Hematoxylin and Eosin (HE), Safranin Red (sGAG), Picrosirius Red (Collagen) and Alizarin Red (Calcium) stains of MSC assembled microtissues. (C) Biochemical quantification of total DNA, sGAG and collagen of MSC assembled microtissues. The data are expressed as mean ± SD. The asterisks indicate p-values obtained by nonpaired two-way ANOVA followed by Tukey's multiple comparisons post-test (*$p < 0.05$; **$p < 0.01$; ***$p < 0.005$; ****$p < 0.001$). Scale bars: (A) - 50 µm and (B) 200 µm.*

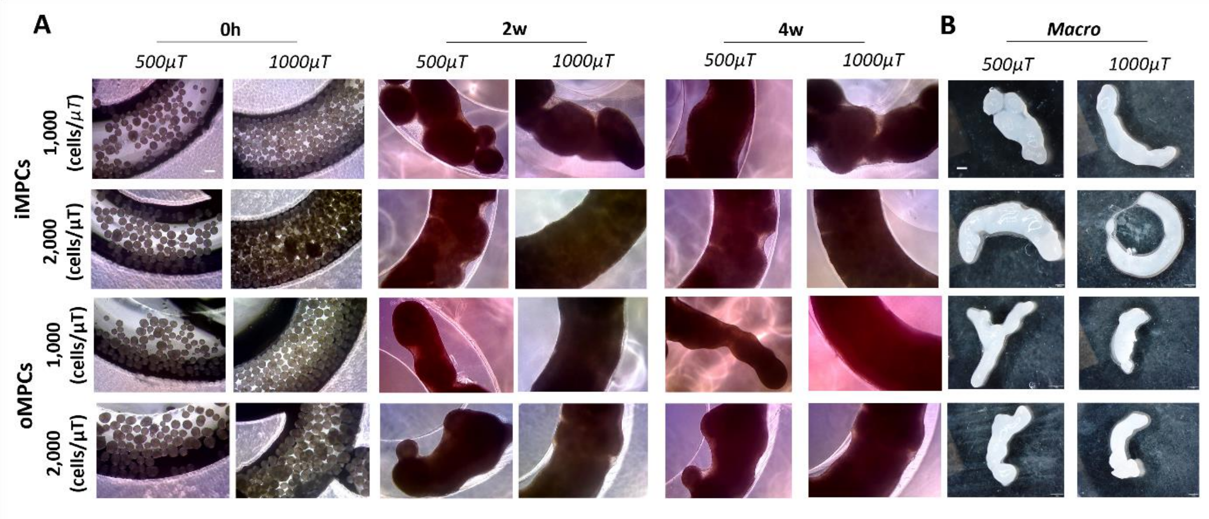


***Supplementary figure 3: Low densities of iMPC and oMPC microtissues do not result in a ring shape tissue.*** *(A) Phase contrast images of approximately 500 and 1000 iMPC and oMPC microtissues fabricated in the densities of 1x10$^3$ and 2x10e$^3$ cells fused in a ring shape mould at 0h, 2 and 4 weeks. (B) Macroscopic images of iMPC and oMPC assembled microtissues. Note that no ring shape was formed after 4 weeks of in vitro culture. Scale bars: (A) – 100 µm; (B) – 500 µm.*

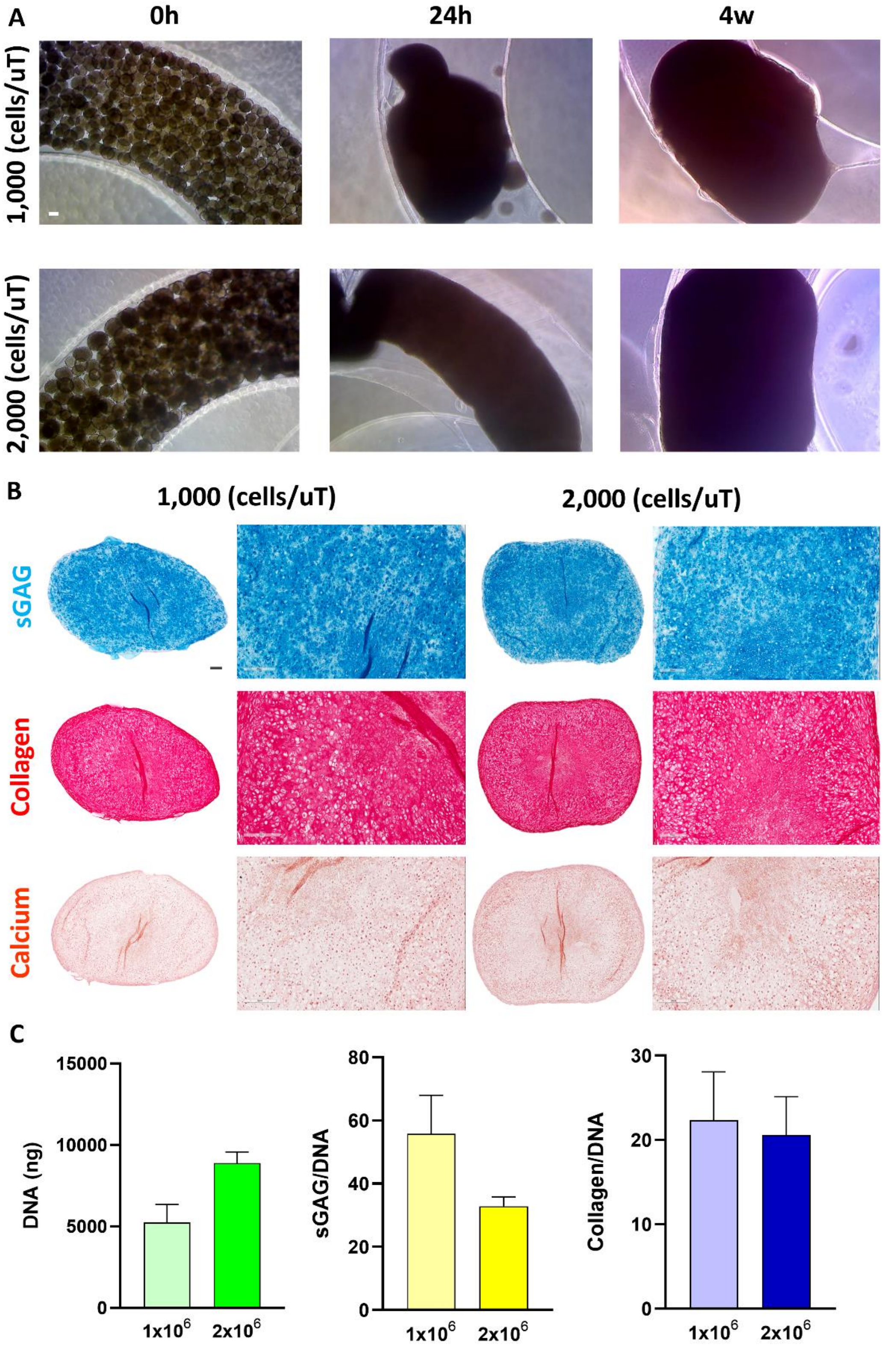


***Supplementary figure 4: MSCs undergo contraction and show lower shape fidelity in a ring shape mould.*** *(A) Phase contrast images of MSC assembled microtissues fabricated using densities of 1x10*$^3$ *and 2x10*$^3$ *cells at 0h, 24h and 4 weeks. (B) Alcian Blue (sGAG), Picrosirius Red (Collagen) and Alizarin Red (Calcium) stains of MSC assembled microtissues at 4 weeks of culture. (C) Biochemical quantification of total DNA, sGAG and collagen of MSC assembled microtissues at 4 weeks of culture. Scale bars: (A) – 100 µm; (B) 500 µm and – 200 µm.*

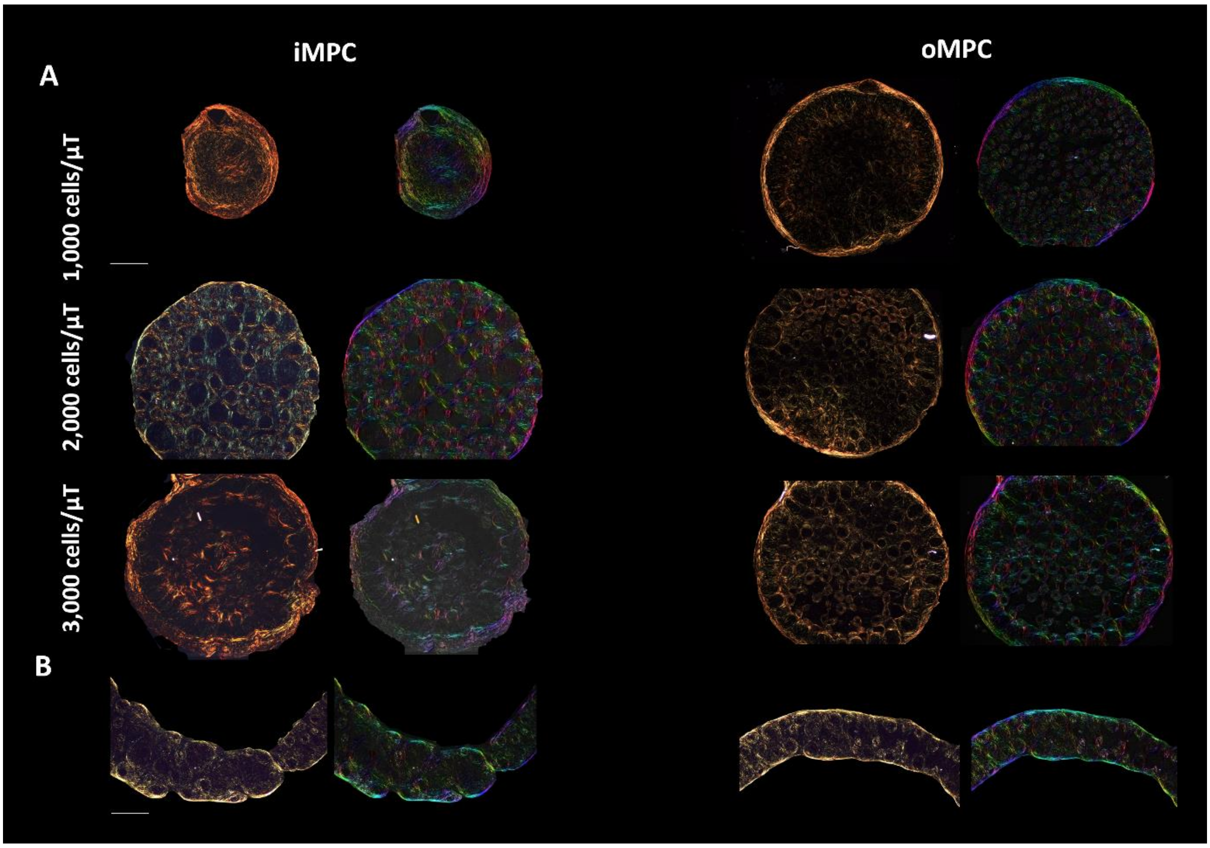


***Supplementary figure 5: Polarized light microscopy (PLM) of iMPC and oMPC microtissues fused in a cylindrical and ring shape mould.*** *(A) PLM of iMPC and oMPC microtissues fused at different densities in a cylindrical shape mould. (B) PLM of iMPC and oMPC microtissues fused in a ring shape mould. Color maps were generated from PLM images. Here, color hue is used to indicate fiber orientation where, red/pink denotes fibers oriented at 90° and blue/cyan indicates fibers are oriented at 0°. Scale bars: (A) – 200 µm and (B) – 400 µm.*

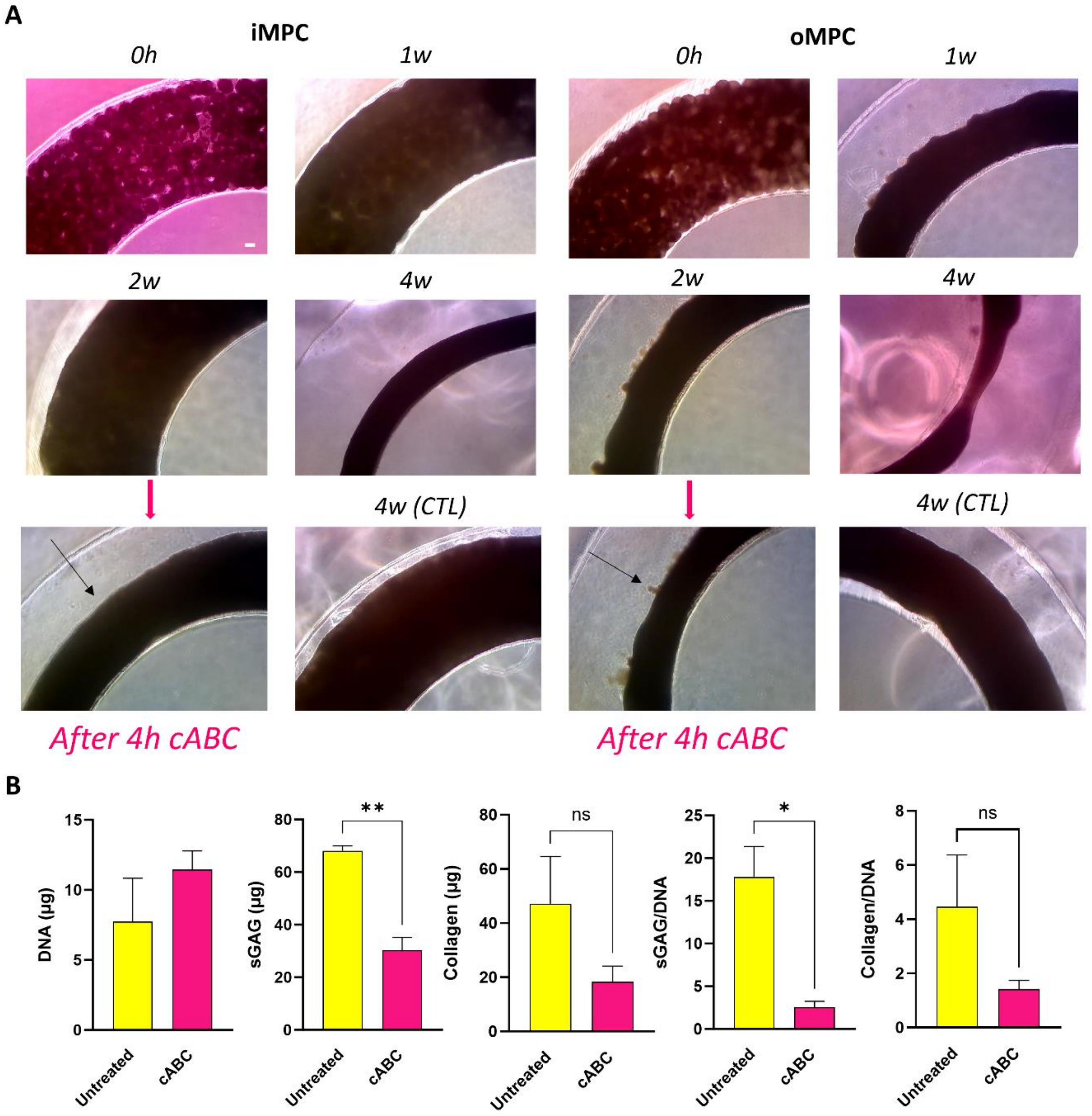


***Supplementary figure 6: Effect of cABC enzymatic treatment in iMPC and oMPC ring assembled microtissues.*** *(A) Phase contrast of iMPC and oMPC ring assembled microtissues at 0h, 1 week, 2 weeks and 4 weeks prior and after 4 hours of enzymatic treatment. Note that iMPC and oMPC ring assembled microtissues had a reduction in size after 4 hours of enzymatic treatment (arrow). (B) Size measurements of iMPC and oMPC ring assembled microtissues prior and 4 hours after enzymatic treatment. (C) Biochemical quantification of total DNA, sGAG and collagen of oMPC ring assembled microtissues 24 prior and after cABC enzymatic treatment. The data are expressed as mean ± SD. The asterisks indicate p-values obtained by nonpaired two-way ANOVA followed by Tukey's multiple comparisons post-test (*p < 0.05; **p < 0.01; ***p < 0.005; ****p < 0.001). Scale bars: 100 µm.*
